# Membrane fusion-based drug delivery liposomes transiently modify the material properties of synthetic and biological membranes

**DOI:** 10.1101/2024.08.14.607934

**Authors:** Jayna Hammond, Ceri J. Richards, YouBeen Ko, Thijs Jonker, Christoffer Åberg, Wouter H. Roos, Rafael B. Lira

## Abstract

Many drug targets are located in intracellular compartments of cells but they often remain inaccessible to standard imaging and therapeutic agents. To aid intracellular delivery, drug carrier nanoparticles have been used to overcome the barrier imposed by the plasma membrane. The carrier must entrap large amounts of cargo, efficiently and quickly deliver the cargo in the cytosol or other intracellular compartments, and must be as inert as possible. In other words, they should not induce cellular responses or alter the cell state in the course of delivery. Here, we show that cationic liposomes with high charge density efficiently fuse with synthetic membranes and the plasma membrane of living cells. Direct fusion efficiently delivers large amounts of cargo to cells and cell-like vesicles within seconds, bypassing slow and often inefficient internalization-based pathways. These effects depend on liposome charge density and, to some extent, liposome concentration and the helper lipid. However, fusion-mediated cargo delivery results in the incorporation of large amounts of foreign lipids that leads to changes in the material properties of these membranes, namely modifications in membrane packing and fluidity, induction of membrane curvature, decrease in surface tension and the formation of (short-lived) pores. Importantly, these effects are transient and liposome removal allows cells to recover their state prior to liposome interaction.

## Introduction

Intracellular delivery of water-soluble cargos using nanosized carriers has long been envisioned as a way to efficiently deliver various biologically-active molecules, nanoparticles and genes into cells. Dependent on their design, nano-carriers can offer controlled and targeted release^[1,2]^, high biocompatibility^[3]^, improved cargo solubility^[4]^, combined imaging and therapeutic modality cargos^[5]^ as well as cargo co-delivery^[6,7]^. Furthermore, they can prevent cargo degradation^[8]^, and enhance cellular uptake^[9,10]^. Among the various delivery vehicles are liposomes: vesicles made of lipids whose fundamental building blocks are similar to the main building blocks of cellular membranes. As delivery systems, they offer large versatility, can encapsulate both hydrophilic and hydrophobic particles at high yield, are easy to produce, purify and functionalize, and their mechanical properties, which often dictate their interaction with the target system^[11]^, can be easily tuned by the choice of their lipid constituents^[12,13]^. An important feature of drug delivery systems is that they are usually designed to be inert, functioning solely as a vehicle for cargo delivery without interfering with any aspect of the target system. However, particles are seldomly completely inert. For *in vivo* delivery, incompatible particles may induce an immune response, potentially leading to adverse side effects. For *in vitro* applications such as cell transfection, it is important that the cells remain alive and functioning for further usage (i.e. ex-vivo transfection and cellular reintroduction). Commonly, nano-biocompatibility is assessed by determining cellular viability, cytotoxicity, or proliferation after exposure to particles^[3]^. However, the effects of nanoparticle carriers on the material properties of the target system has been generally overlooked.

In general, nanoparticles enter cells via endocytosis, or in some instances, via direct penetration through the cell membrane^[14,15]^. However, both delivery methods have their drawbacks ^[16]^. In the case of endocytosis, the particles are typically trafficked through the endo-lysosomal pathway, introducing the additional challenge of releasing the drug from the endo/lysosomes into the cell cytosol before degradation or exocytosis^[14,15]^. Particles that enter through direct penetration, on the other hand, may have off-site delivery and may induce damage to the cell plasma membrane (PM)^[17,18]^. In contrast, a specific class of liposomes that deliver their cargo via fusion with the PM, called fusogenic liposomes, are able to directly deliver cargos into the cell cytosol whilst maintaining targeting capacity^[19,20]^. The direct fusion of liposomes inevitably involves the transfer of the lipid content from the liposome to the cell PM. This potentially induces changes in the properties of the PM, either locally at the fusion site, or more globally across the entire cell. Which and to what degree the PM material properties change due to membrane mixing in the course of drug delivery, whether the effects are adverse, and finally if they are reversible are important questions that have remained elusive. Moreover, for *in vivo* nano carrier applications, the adsorption of environmental proteins onto the carrier surface, known as biomolecular corona formation^[21]^, has been shown to mitigate toxicity in many systems^[22]^, although in some cases it may reduce targeting ability^[23,24]^ or prevent delivery altogether^[25–27]^. Thus, the effect of environmental proteins on the fusion process, and hence on intracellular delivery of cargos, and any induced cellular changes are important factors to consider, but remain largely unexplored.

Here, we set out to investigate whether liposome nanovehicles, containing or devoid of cargo, pose short- or long-term effects on acceptor synthetic and biological membranes. For this, we use cationic fusogenic liposomes that readily (thousands per cell) and quickly (within seconds) fuse with acceptor membranes, efficiently delivering their encapsulated cargo. To assess the specific effects of charge or chemical composition of the lipid formulation, we use either mono or multivalent cationic lipids with varying liposome charge density, containing cylindrical or cone-shaped helper lipids. We used multicolour confocal microscopy and fluorescence lifetime imaging microscopy (FLIM) to assess fusion efficiency and the effects of fusion on the material properties of membranes, including changes of membrane fluidity, membrane remodelling and formation of leakage pores. As model systems, we used giant unilamellar vesicles (GUVs) that are devoid of metabolism, receptors or other active processes, so the physical-chemical effects on membranes are assessed without interference from underlying biological activity, as well as living cells to understand how these changes impart on biological systems.

We show that cationic liposomes exhibiting a high charge density (σ_M_), efficiently fuse with acceptor synthetic and biological membranes, efficiently delivering cargo. At this σ_M_, helper lipids become dispensable. At intermediate σ_M_, fusion efficiency is reduced despite stable membrane docking, and incorporation of inverted cone-shaped helper lipids improves fusion efficiency. Uncharged liposomes are unable to bind and fuse. Interestingly, pre-incubation in serum leads to protein corona formation on the liposome surface, shielding their charges and having a similar effect as low σ_M_, preventing fusion. Extensive liposome fusion modifies acceptor membrane fluidity in a way that can be predicted by lipid compositional changes, leading to changes in membrane tension, curvature and eventually to the formation of leakage pores. Interestingly, the formed pores have specific sizes and a limited lifetime. This means that even if the PM of living cells become permeabilized, the delivered cargos are retained in the cytosol. Importantly, removal of liposomes from the media recovers the introduced changes in the membrane material properties.

### 1. Rationale

In our proto drug delivery system, strong interaction and fusion with acceptor synthetic and biological membranes dispersed in low ionic strength solutions requires charge. To enable a strong interaction, and thereby a higher probability of fusion between the liposome and acceptor membranes, we decided to use positively charged liposomes. Admittedly, electrostatic interactions are highly screened in high ionic strength solutions such as serum, but the interaction should nevertheless remain strong at short range. Furthermore, ex vivo experiments can readily be performed under low ionic (but isotonic) conditions, as done here, where the electrostatic interactions will remain strong. To render the liposomes positively charged, their membranes contained the cationic lipid DOTAP. DOTAP is a cylindrically shaped lipid that forms bilayers, and hence liposomes can be made of purely DOTAP lipids. Furthermore, it is well known that the structure and activity of liposomes also depends on helper lipids, conferring liposome stability, determining phase state and restructuring upon interaction with target membranes^[28,29]^. The most commonly used helper lipids are DOPC and DOPE. Due to its cylindrical shape, DOPC forms lamellar phase^[30]^, and thus liposomes can also be made purely of DOPC. In contrast, DOPE is an inverted cone-shape lipid that imposes negative membrane curvature, and DOPE alone or with low fractions of cylindrical lipids forms inverted hexagonal phases^[31]^. Depending on the helper lipid, liposomes exhibit different modes of interactions with cells, with mixtures of DOTAP:DOPE efficiently fusing with the cell plasma membrane, whereas DOTAP:DOPC liposomes are internalized by cells^[32,33]^.

To separately determine the roles of membrane charge and of the helper lipid on the interaction and fusion of liposomes, we used giant unilamellar vesicles (GUVs) as a model of the PM. The GUVs were composed of equimolar mixtures of DOPC:PS:Chol:DOPE, in which PS stands for brain phosphatidylserine, (or alternatively, DOPG was used instead of PS without measurable differences), as this composition closely mimics some mechanical properties of the PM, including fluidity and charge^[34,35]^. Interaction was studied with the recently developed time-resolved fluorescence resonance energy transfer assay^[36]^. In this assay, fluorescence resonance energy transfer (FRET) is assessed with fluorescence lifetime imaging microscopy (FLIM). GUVs are doped with the donor Bodipy C_16_ probe and the liposomes are made with the acceptor lipid Rh-PE. Bodipy C_16_ is nearly insensitive to the environment^[37]^ and thus lifetime changes are only associated with FRET. Fusion of liposomes with the GUVs results in an increase in donor-acceptor proximity and a reduction in donor lifetime due to FRET, whereas binding only causes minimal FRET changes^[38]^. To resolve if the liposomes fully fused with the GUVs, the liposomes encapsulated the water-soluble dye sulforhodamine B (SRB). Upon complete merging (full fusion) of the GUV and liposomal membranes, SRB is transferred to the GUV lumen. On the other hand, the appearance of SRB-containing liposomes on the GUV surface without content mixing is an indication of liposome binding without fusion. Therefore, in this configuration, FLIM-FRET is able to resolve membrane docking, hemifusion and full-fusion at the single vesicle level (Figure 1).

**Figure 1.**
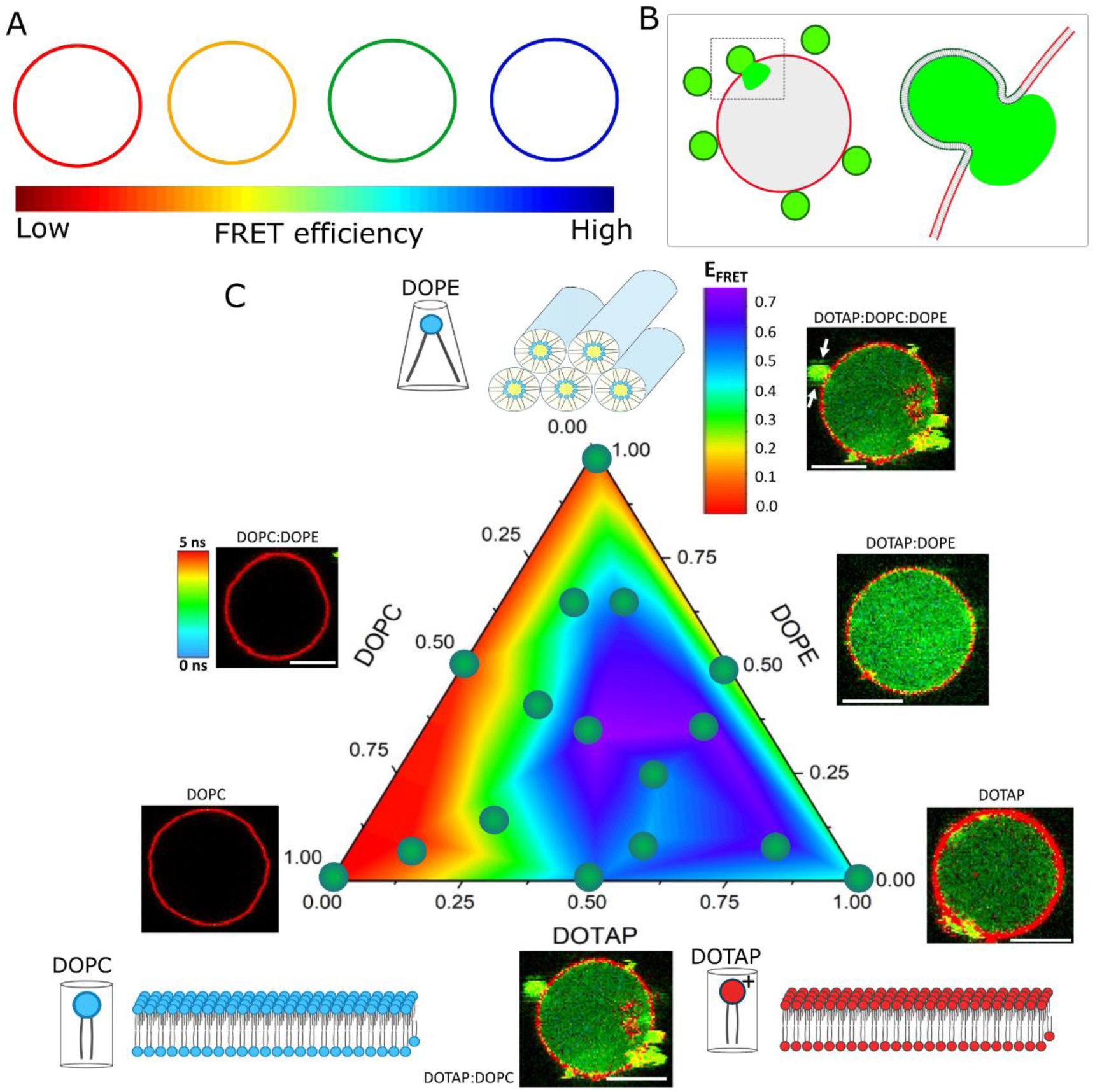
Fusion efficiency diagram for three-component liposomes. A, FLIM-FRET sketch of the fusion assay. Donor lifetime decreases (red to blue colours) as fusion efficiency increases, as assessed from an increase in FRET efficiency (E_FRET_). B, sketch of the content mixing assay in which liposomes encapsulating a water-soluble probe (green) fully fuse with an acceptor GUV (red), transferring its content to the GUV lumen. C, E_FRET_ fusion diagram measured on GUVs incubated with liposomes containing varying fractions of DOPC, DOPE and DOTAP. The sketches show that DOPC alone forms neutral bilayers (lower left), DOTAP alone forms positively charged bilayers (lower right), whereas DOPE alone forms an inverted hexagonal phase (top) due to its molecular geometry. The heat map within the ternary component diagram represents the estimated fusion efficiency based on E_FRET_ measurements. Each green solid circle represents a tested liposomal composition (10-15 GUVs per liposome composition). Insets: GUV FLIM images show representative content mixing results from incubation of GUVs with the respective liposomal compositions. Here, unlabelled liposomes encapsulating 50 μM SRB fuse with Bodipy C_16_-containing GUVs. Bodipy C_16_ and SRB are clearly resolved in the FLIM images due to their lifetime differences. Note that in some of the images, docked liposomes can be observed (arrows). Scale bar: 8 μm. The GUVs are made of DOPC:PS:Chol:DOPE (25:25:25:25, mole fraction).

### 2. Fusion efficiency is determined by membrane charge and modulated by helper lipids

To independently assess the effects of membrane charge and cylindrical or inverted cone-shaped helper lipids, we varied the mole fraction of DOPC, DOTAP and DOPE in the liposomes. To assess fusion efficiency, we measured FRET efficiency (E_FRET_) at the single GUV level. In this manner, we were able to construct a liposomes composition fusion diagram for which FRET is used as the readout of fusion efficiency (Figure 1). We started with the single component compositions. As expected, liposomes made of the neutral DOPC alone do not fuse with the GUVs (E_FRET_ ∼ 0). DOPE alone does not form bilayers, and the hexagonal phase lipid aggregates mixed with the GUVs also do not induce any change in FRET. In contrast, liposomes made purely of DOTAP efficiently fuse. These results show that charge is necessary and sufficient for fusion. We next examined two-component mixtures of lipids. As expected, neutral liposomes composed of DOPC:DOPE do not undergo fusion. In contrast, DOTAP mixtures with either DOPC and DOPE efficiently fuse with the GUVs. Interestingly, while DOPC does not seem to change the efficiency of fusion of DOTAP, mixtures with DOPE result in higher E_FRET_ compared to that of DOTAP alone. Of note, this binary composition is more efficient than any ternary mixtures of these lipids, indicating that the cylindrical DOPC lipid reduces liposome fusion efficiency. Aside from the results shown in Figure 1 C, representative lipid mixing FLIM GUV images are shown in Figure S1 along with their respective donor decays. From the fusion diagram, it is clear that liposomal compositions rich in DOTAP and DOPE are the most fusogenic, whereas compositions further away from this mixture exhibit lower fusion efficiencies.

Importantly, for all compositions that exhibit high E_FRET_, the interaction results in the transfer of SRB from the liposomes to the GUV interior, showing that these liposomes undergo full fusion (inset GUV images in Figure 1). Liposome docking can also be resolved from diffraction-limited SRB-containing liposomes (green) on the GUV surface. In contrast, the compositions that do not lead to changes in E_FRET_ also do not lead to full fusion. In fact, liposomes made of neutral lipids alone do not bind the GUVs, indicating that they are unable to fuse because they are unable to bind in the first place. In summary, the results show that inclusion of the cationic lipid DOTAP in the liposome is required and sufficient to efficiently fuse liposomes with biomimetic membrane and the addition of the inverted cone-shape helper lipid DOPE to the liposome further enhances fusion.

### 3. Charge density, not chemical identity, determines membrane fusion efficiency and fusion-dependent pore formation

The maximum σ_M_ attained with the liposomes using DOTAP as the cationic lipid (CL) is when DOTAP mole fraction (ϕ_DOTAP_) is 100%, thus devoid of helper lipids. However, higher σ_M_ could be attained with multivalent CLs at lower mole fractions, with the additional advantage that helper lipids could be added for fine tuning liposome properties. Furthermore, it has been shown that cationic liposomes can be toxic to cells^[39,40]^ and have limited applicability in physiological media containing serum proteins due to charge screening and particle aggregation^[41]^. In this section, we study the effects of liposome charge density on fusion with GUVs. For this, we increase the mole fractions of the monovalent lipid DOTAP and multivalent CL lipid MVL5 (ϕ_MVL5_), a lipid that harbours up to 5 positive charges depending on pH. For these formulations, we also investigated the effects of cylindrical (DOPC) or negatively-curved (DOPE) helper lipids. Later, we assessed whether liposome fusion modifies the structure of the acceptor GUV membrane (i.e. formation of pores) and how interaction with serum proteins affects these processes.

We prepared liposomes with increasing mole fractions of the CL (ϕ_CL_) DOTAP or MVL5, containing DOPC alone or DOPC:DOPE mixtures as the helper lipid(s) (Figure 2 A). These compositions are represented as CL:DOPC (X:100-X) or CL:DOPC:DOPE (X:50-X:50), in which X is the fraction of the CL, adjusted at the expense of DOPC. Figure 2 B shows representative FLIM images of GUVs incubated with CL:DOPC:DOPE liposomes with increasing ϕ_CL_. Note that as ϕ_CL_ increases E_FRET_ also increases (Figure 2 C). The quantitative E_FRET_ response curves as a function of ϕ_CL_ in the membrane for liposomes containing DOPE or not, are shown respectively in Figures 2 D and 2 E. At identical ϕ_CL_, E_FRET_ is much more pronounced with MVL5. At very low ϕ_CL_, liposomes containing DOTAP are not fusogenic, whereas fusion is readily detected with as low as 1 mol% MLV5. The fraction at which fusion is half maximum is ∼ 12 or 28 mol%, for DOTAP liposomes with or without DOPE, respectively, again pointing to a modulating effect of DOPE on fusion, and 1.3 mol% for MVL5 irrespective of the helper lipid. For both CL (or helper lipids), the maximum efficiency saturates. However, the saturation occurs at much lower fractions of MVL5 (∼ 2.5%) compared to DOTAP (∼ 25%). Thus, much lower amounts of multivalent CL is required to achieve maximal fusion.

**Figure 2.**
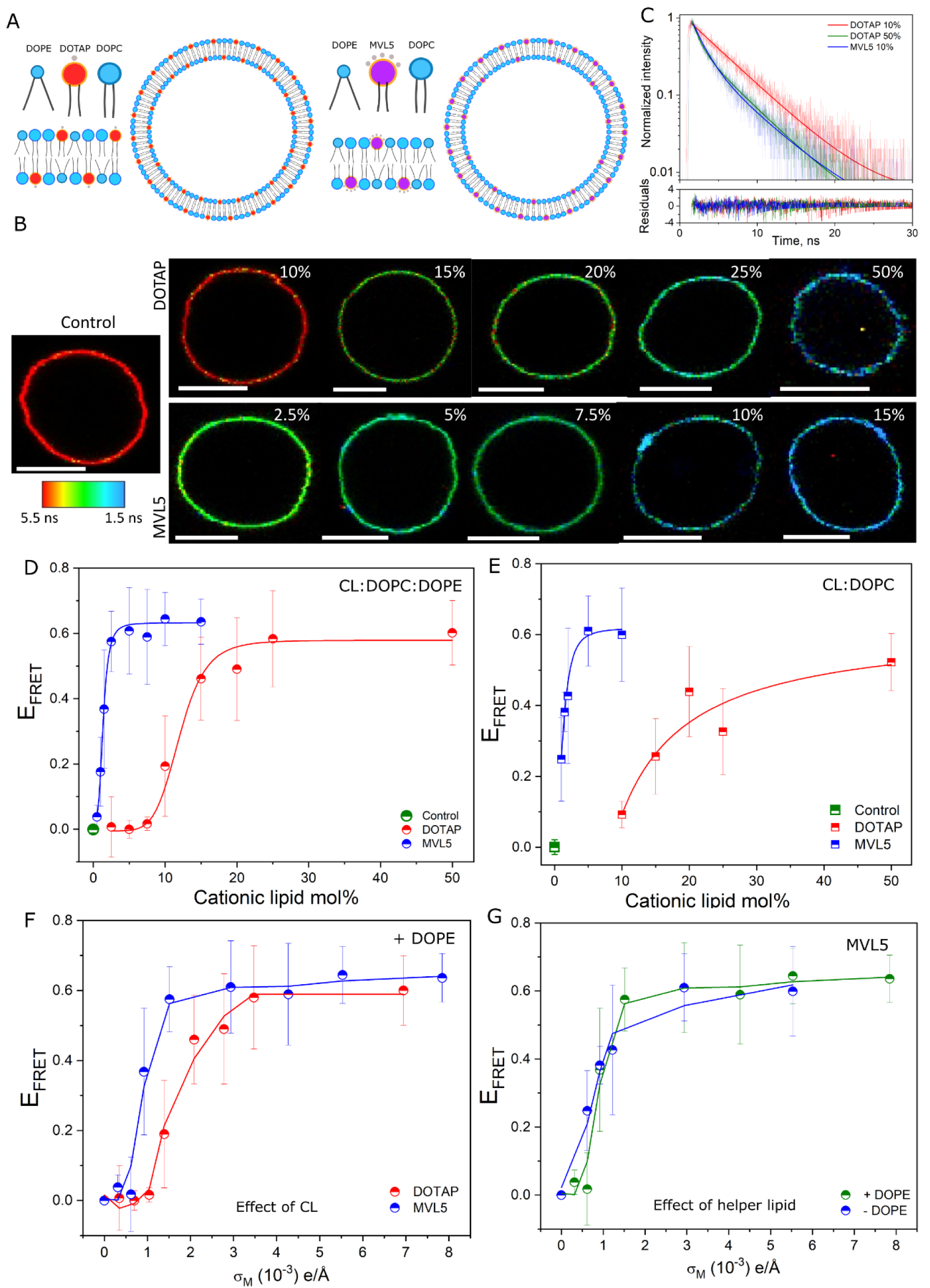
Membrane charge density is the main factor determining fusion with acceptor membranes. A, sketch of liposomes containing the monovalent lipid DOTAP or the multivalent lipid MVL5 in the presence of helper lipids DOPC and DOPE. B, representative FLIM GUV DOPC:PS:Chol:DOPE (25:25:25:25, mole fraction) images upon incubation with CL:DOPC:DOPE (X:50-X:50, mole fraction) 20 μM liposomes (lipid concentration) containing increasing mol% of the CL and labelled with 2 mol% DPPE-Rhodamine. Control GUVs in the absence of liposomes are also shown along with the lifetime map. Scale bar: 8 μm. C, representative lifetime decays, respective fits and residuals from the fits for GUVs labelled with FRET donor Bodipy C_16_ upon fusion with several liposome compositions. D and E, fusion efficiency as represented by E_FRET_ as a function of CL mole fraction for liposomes containing DOPE and DOPC or only DOPC as the helper lipid, respectively. Control represents measurements in the absence of liposomes. Each data point represents measurements on 10-15 individual GUVs. Means and s.d. are shown. The lines are a guide to the eye. F, E_FRET_ as a function of the calculated charge density for liposomes containing DOTAP or MVL5 and DOPC:DOPE as the helper lipids. Liposome fusion efficiency strongly depends on σ_M_ and weakly on the chemical identity of the CL. G, E_FRET_ response as a function of charge density for liposomes either containing DOPE or not as the helper lipid and MVL5 as the CL. The response does not have a strong dependence on the helper lipid.

At maximum fusion, the ϕ_DOTAP_ is approximately 10-fold higher than ϕ_MVL5_ despite the < 5 charge ratio between them. To isolate the effects of charge, we replotted the data as a function of σ_M_, which is given by^[42]^

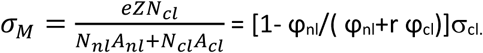

Here, N_nl_ and N_cl_ are respectively the number of neutral and cationic lipids; Z is the valence of the CL (we used 1 and 4.5 for DOTAP and MVL5, respectively), r = A_cl_/A_nl_ is the area ratio of the cationic and neutral headgroups (we used 2.3 here, with A_nl_ being 72 Å^2^ ^[42]^), and σ_CL_ is the mole fraction of the cationic lipid. The replotted E_FRET_ versus the calculated σ_M_ response nearly falls onto a universal curve for DOPC only (Figure S2 A) or with DOPC:DOPE mixtures as helper lipids (Figure 2 F), irrespective of the CL. Thus, σ_M_ seems to be the determining factor for fusion, not the specific chemical identity of the CL. Therein, the role of the helper lipid is secondary to σ_M_, with comparable E_FRET_ changes for DOTAP or MVL5 liposomes (Figure 2 G and S2 B, respectively). At low σ_M_ (< 10^-3^ e/ Å), fusion is negligible, whereas fusion is equally efficient at high σ_M_ (> 3-4 10^-3^ e/Å) for both CLs, with a mildly higher efficiency for liposomes containing DOPE, as expected from the fusion diagram in Figure 1. Importantly, in all cases with DOTAP or MVL5, and for liposomes containing either DOPE or not as the helper lipid, the interaction with GUVs is via full fusion (Figure S3). The results show that it is σ_M_ and not the chemical identity of the CL that is the determining factor for fusion of cationic liposomes with acceptor membranes, with efficient delivery of cargo into GUV model membranes, and with the helper lipids having a regulatory effect on fusion at moderate σ_M_.

A disadvantage of using cationic nano(particles) as delivery vehicles is that they are known to be toxic to cells^[18,39,40,43]^. However, it is not known whether the toxic effects are associated with the total charge (i.e. σ_M_) that needs to be processed by cells, in which case DOTAP and MVL5 would be equally toxic, or with the number of charged molecules that need to be dealt with, in which case DOTAP would be more toxic than MVL5 at identical σ_M_. Understanding the molecular mechanisms of toxicity will help the design of efficient drug delivery vehicles with limited toxicity. As a proxy of toxicity, we investigated the effect of liposome fusion on pore formation in GUVs. Recently, we have shown that extensive fusion may lead to the formation of hydrophilic pores in GUV membranes, an effect that is suppressed by physiological levels of cholesterol^[36]^. We labelled the liposome membranes with Atto647-PE (cyan) as a fusion reporter, and co-incubated these liposomes with GUVs in the presence of sulforhodamine B in the medium (SRB, red), a water-soluble probe the can only transverse membranes through pores. The assay is able to simultaneously detect fusion, fusion-dependent pore formation, as well as vesicle morphology. We use the degree of filling analysis to quantitatively assess the extent of disruption^[44]^ wherein values close to unity indicate that larger or longer-lived pores (or a combination of both) are formed, a sign of more extensive disruption.

We reasoned that more fusion would result in more disruption, in which case more intense membrane signal would be accompanied by more SRB leakage into the GUVs (Figure 3 A). Figure 3 B shows two representative GUVs incubated with DOTAP-containing liposomes (ϕ_DOTAP_ 0.5) that either do or do not contain DOPE as the helper lipid. As anticipated, more extensive fusion (stronger membrane signal) results in leakage (SRB entry). Figures 3 C and D show the effects of liposome fusion as a function of ϕ_CL_ for DOPC or DOPC:DOPE helper lipid(s). As expected, fusion becomes more efficient as ϕ_CL_ increases. Importantly, in general, the process does not lead to extensive pore formation. The only exception is the highly charged liposomes containing 50 mol% DOTAP but without DOPE. For those vesicles that do become permeable, the SRB signal in the GUV interior is lower than that in the outside medium, an indication that the formed pores reseal before full SRB equilibration. Although in general the permeabilization effects are mild, more disruptive effects such as complete GUV collapse can also be observed (Figure S4 and Movie S1)^[45]^. Of note, in addition to ϕ_CL_, fusion efficiency also depends on liposomal concentration (Figure S5). When we plot the degree of filling for increasing σ_M_, the GUVs only become permeable at high σ_M_, (Figures 3 E and F). However, this effect again does not depend on the CL lipid used, although the presence of DOPE decreases the fraction of permeable GUVs. The results show that, similarly to the effects on fusion, the charge density, not the chemical identity of the CL, is the determining factor for fusion-dependent membrane disruption, with the helper lipid having a tuning effect on the process. In this case, DOPE tends to both increase fusion efficiency and decrease fusion-dependent disruption.

**Figure 3.**
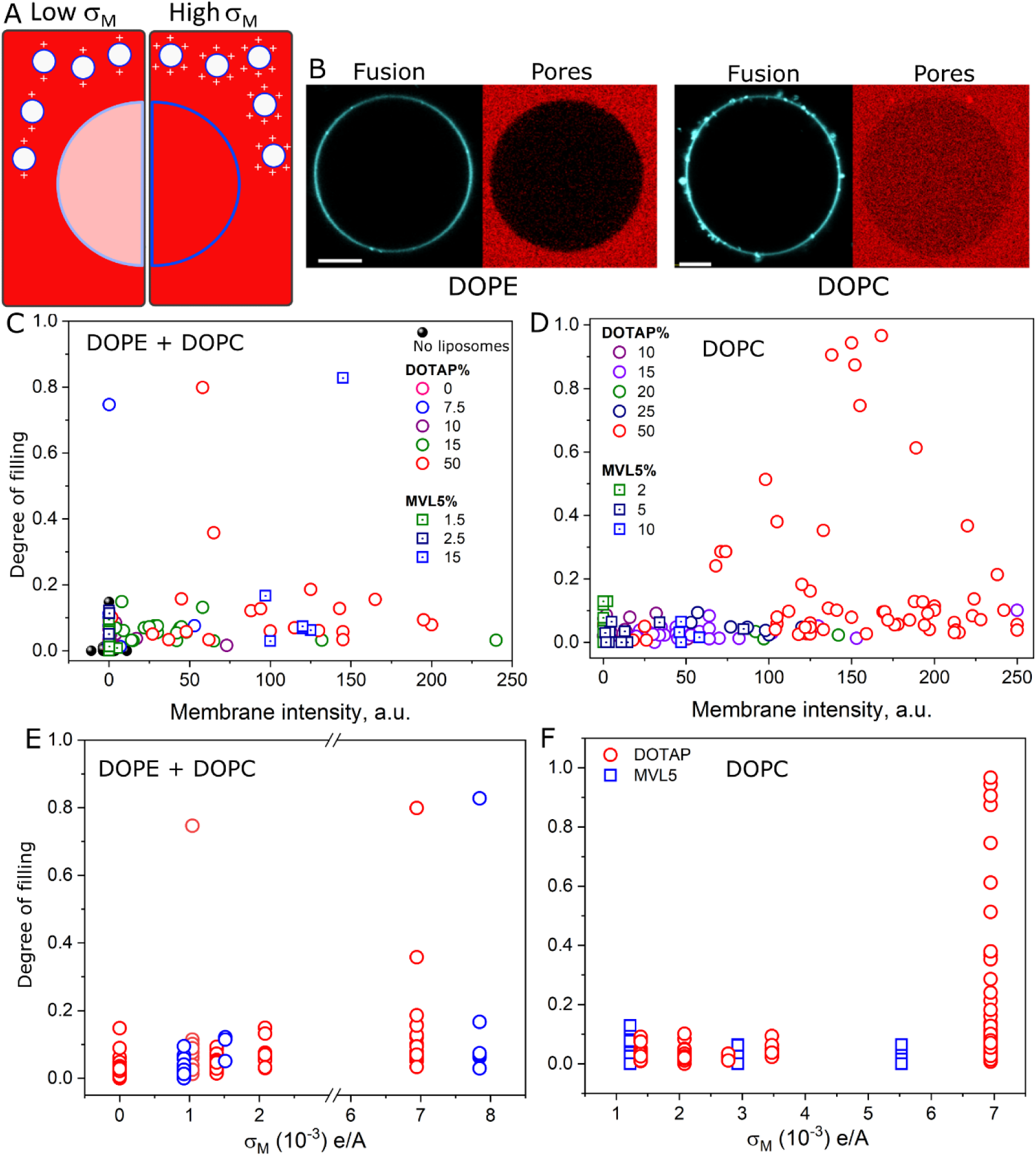
Charge density determines fusion efficiency and membrane disruption. A, sketch of the fusion/permeabilization assay with low (left) and high (right) charge density (σ_M_) liposomes labelled at the membrane with Atto647-DOPE (cyan) and incubated with GUVs (DOPC:DOPG:Chol:DOPE, 25:25:25:25, mole fraction) in the presence of 10 mM SRB (red) in the outer solution used as a leakage marker. More fusion (more intense membrane signal) is expected to result in more membrane permeabilization (more SRB entry). B, representative GUV images incubated with DOTAP (50 mol%) liposomes (20 μM total lipids) either containing DOPE as the helper lipid or not (DOTAP:DOPE or DOTAP:DOPC, respectively). Scale bars: 10 μm. Note that higher fusion (more intense cyan signal at the membrane) co-occurs with more SRB entry into the GUVs. C and D, degree of filling dependence on fusion efficiency (measured as the intensity of Atto647-DOPE in the GUV membrane) for DOTAP or MVL5 as the CL in liposomes containing DOPE and DOPC or only DOPC as the helper lipid, respectively. Liposome composition: CL:DOPC:DOPE (X:50-X:50, mole fraction). The control without incubation of liposomes is also shown (no liposomes). E and F, degree of filling in GUVs incubated with liposomes with increasing charge densities (σ_M_) for liposomes containing or not containing DOPE as the helper lipid. Each point represents a measurement on a single GUV.

### 4. Fusion modifies membrane fluidity and curvature

Membrane fusion results in the mixing of two initially separated membranes. Thus, unless these membranes have identical compositions, fusion inevitably changes their lipid make up. Membrane physical properties are strongly tied to composition^[12]^. Since the compositions of the liposomes and GUVs used here are not identical, we speculate that liposome fusion will not only result in changes in GUV composition, but also in their mechanics. In fact, we have previously shown that full fusion of DOTAP:DOPE liposomes to POPC:POPG GUVs induced a clear reduction in lipid diffusion in these GUVs, whereas only a milder decrease in diffusion was observed when the liposomes hemifused to neutral POPC GUVs^[38]^. Using the Saffman-Delbrück equation^[46]^ (see methods), we calculated the approximate membrane viscosity in these membranes from the reported lipid diffusion values. The obtained values are comparable to reported FLIM data using viscosity-sensitive molecular rotors^[37]^. As shown in Figure S6, fusion increases the viscosity of POPC:POPG GUVs. The composition dependence is confirmed since mimetic GUVs – vesicles made of the expected composition after fusion – also exhibited increased viscosity. It has been previously reported that, in the absence of other factors, DOPE increases membrane viscosity^[47]^. We thus interpreted these results as a consequence of fusion-mediated incorporation of DOPE in the GUV membranes.

To confirm these observations, we used FLIM combined with molecular flippers, molecules that undergo excited-state changes in their intensity and lifetime as a function of environmental packing^[48,49]^, as a more direct way to assess membrane packing, which relates to viscosity. As observed previously, FliptR’s fluorescence decay is best fitted with a double exponential decay, in which the long lifetime (τ_long_) reports packing^[48,49]^. Figure S7 shows that liposome fusion with fluid DOPC:DOPG GUVs leads to an increase in τ_long_ (thus an increase in packing) and this increase is weakly dependent of the CL or helper lipid. To test the generality of these observations, we studied fusion of liposomes with initially more ordered membranes rich in Chol. If the viscosity/composition dependence holds, liposome fusion would then result in a decrease rather than increase in viscosity. For this, we prepared GUV membrane models containing (i) highly charged and fluid membranes, and (ii) low charge and highly packed membranes, both rich in cholesterol. These two compositions mimic the lipid compositions of the inner and outer leaflet of the PM, respectively, and we have recently shown that the combination of charge and packing allows efficient fusion with the former, but not with the latter^[36]^. Figures S8 A shows representative GUVs from these compositions before and after liposome fusion, as well as mimetic GUVs with the expected composition upon complete charge neutralization. Figure S8 B shows the respective fluorescence decay. As expected, fusion of liposomes with GUVs of composition (i) indeed decrease τ_long_, showing a reduction in membrane packing (the lifetime ratio of liposome-incubated/control ∼ 0.95), whereas liposomes interacting with GUVs of composition (ii) show detectable but milder effects (∼ 0.98 ratio) due to the liposome reduced ability to fuse with these membranes.

In addition to the changes in fluidity, liposome fusion also modifies the spontaneous curvature of the GUVs. As can be seen in figure 3 B and Figure S9, the GUVs that underwent extensive fusion (strong membrane signal) also exhibited outward buds. This effect has been observed previously and is interpreted as a result of the transfer of the leaflet asymmetry number, inherent to small liposomes, to the otherwise flat membranes of the GUVs^[38]^. Since membrane curvature is a physical process and occurs whenever a sufficiently large number of (small) liposomes fuse to the GUVs, bud formation should not depend on the particular liposome composition, as indeed verified (Figure S9). The diameter of the bud reflects the extent of curvature – higher curvatures form smaller buds^[50,51]^. In our experiments, for DOTAP or MVL5 liposomes at identical σ_M_, either containing DOPE or not as the helper lipid, and for all concentrations tested, bud diameter does not vary significantly, as expected given their comparable fusion efficiencies. On the other hand, much higher concentrations result in complete vesicle disruption leading to GUV bursting as in Figure S4, and are thus beyond experimental range. FLIM images of FliptR-labelled GUVs that contain buds show no visible lifetime differences in the buds compared to the flatter regions of the GUVs, indicating composition equilibration, although fitting the data from the buds alone was not possible due to the low photon count. In summary, the results show that the fluidity of the fusing membranes changes as a consequence of the changes in membrane composition, fusion transfers the intrinsic liposome leaflet number asymmetry to the GUVs, changing membrane spontaneous curvature, and lipids are fairly equally distributed in the regions of higher or low curvature.

### 5. Protein corona screens liposome charge and prevents membrane interaction and fusion

In biological systems, media are rich in ionic species as well as large concentrations of biomacromolecules, both of which have the ability to reduce interactions, by charge screening and protein corona formation, respectively^[43,52]^. While membrane fusion of cationic liposomes and GUVs is efficient at high ionic strength^[38]^, here we tested the effects of protein corona formation. We first pre-incubated the cationic liposomes with serum, and then delivered the liposome/serum dispersion to the GUVs. Thus, note that GUVs were exposed to the liposomes in the presence of both a hard and soft corona, as well as in the presence of serum proteins in the medium. Zeta potential measurements on pristine cationic liposomes show comparatively high values > 40 mV for DOTAP and MVL5 (Figure S10). Incubation with serum not only neutralizes the liposome charge, but actually “overshoots” the neutralization such that the zeta potentials become negative, most likely due to dense protein coating on the particle surface. At the serum concentration tested, there are no signs of liposome aggregation irrespective of the CL used (Figure S11). We used multicolour confocal and FLIM microscopy to study the effects of protein corona formation on fusion. Figure 4 A shows representative FLIM images and Figure 4 B shows the fluorescence decay of the FRET donor Bodipy C_16_ for: a control GUV in the absence of liposomes, a GUV incubated with pristine liposomes, and a GUV incubated with liposomes pre-incubated with serum. As expected, pristine liposomes efficiently fuse with the GUVs, reducing the acceptor lifetime compared to the control GUV as a result of fusion. In sharp contrast, liposomes pre-incubated with serum do not undergo fusion as observed by the unmodified donor lifetime. Quantitative analysis on many individual GUVs shows that, for liposomes with comparable σ_M_, fusion is blocked if the liposomes are pre-incubated with serum, irrespective of the CL used (Figure 4C). Similar results are observed for the intensity analysis (Figure S12) and they are not dependent on GUV size (Figure S13).

**Figure 4.**
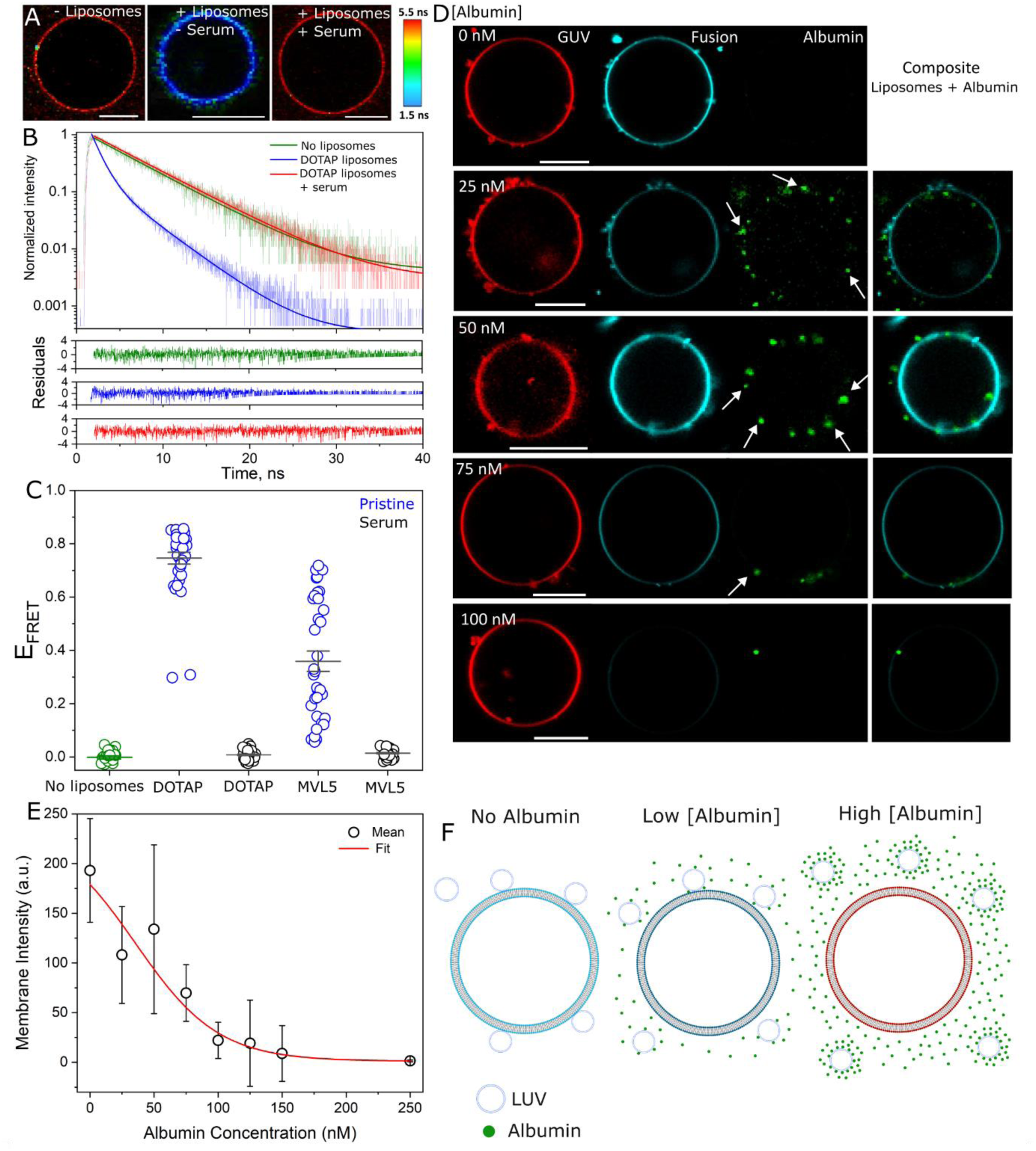
Protein-coated cationic liposomes are unable to fuse with GUVs. A, representative FLIM images of Bodipy C_16_-labelled GUVs (DOPC:DOPG, 1:1 mole fraction) in the absence of liposomes, in the presence of pristine liposomes and in the presence of liposomes pre-incubated with serum (10x diluted). Scale bars: 9 μm. B, Bodipy C_16_ fluorescence decays along with the fits and residuals of the fits for the GUVs shown in panel A. The data from the control (- liposomes) and the liposomes pre-incubated in serum (+liposomes/+serum) were well-fitted by a single-exponential decay, and the data for pristine liposomes (+liposomes/-serum) were well-fitted by a double-exponential decay. C, measured E_FRET_ for a number of GUVs upon incubation with pristine or protein-coated liposomes containing DOTAP (DOTAP:DOPE, 33:66 mole fraction) or MVL5 (MVL5:DOPC:DOPE, 5:50:45 mole fraction) as the CL. The control without liposomes is also shown. D, representative GUVs (labelled with 0.5 mol% DPPE-Rhodamine, red) upon incubation with 20 μM (lipid concentration) Atto647-DOPE (cyan, representing fusion) liposomes that have been pre-incubated with increasing concentrations of FITC-labelled albumin (green). The arrows point to protein particles or complexes with liposomes on the GUV surface The overlay composite of liposome and albumin channels is also shown. Scale bar: 10 μm. E, membrane signal measured from the average Atto647-DOPE fluorescence intensity upon liposome fusion for a number of GUVs (10-15 per condition) as a function of albumin-FITC concentration. Mean and standard deviation are shown. A fit of an exponential decay (Y=A_1_e^(-X/t1)^+y_0_, where A_1_ and y_0_ are the maximum Atto647-DOPE intensity and offset at 0 intensity, respectively) to the data is also shown. F, sketch of the expected interactions between cationic liposomes and GUVs. In the absence of albumin, the liposomes directly fuse with the GUVs, transferring their (cyan) lipids. At intermediate albumin concentrations, partial liposome coverage reduces liposome binding but those that are bound can still fuse. At high protein concentration, complete protein coverage prevents liposome binding altogether and there is no lipid mixing.

The FLIM-FRET assay above precisely detects fusion efficiency but it is insensitive to liposome binding to GUVs. To gain insights into the effect of corona on the separate processes of liposome binding and fusion, we performed multicolour microscopy of liposomes (cyan), a serum model comprised of fluorescently-labelled albumin (green) of well-defined concentrations, and GUVs (red). Albumin is a negatively charged protein and incubation with the cationic liposomes is expected to result in liposome-albumin complex/corona formation. Because GUV imaging was performed in the sequential mode to minimize spectral cross-talk, the slow imaging combined with fast diffusion makes it difficult to assign co-localization between albumin and liposomes. Nonetheless, co-localized green-cyan spots are observed for various albumin concentrations (Figure S14), indicating protein-liposome complex formation. Protein complex binding to the GUV surface is also observed (Figure 4 D). Increasing the concentration of albumin gradually decreases fusion as assessed from the reduced transfer of lipids from the liposomes to the GUVs (Figure 4 D and 4 E). While liposomes in the absence of albumin exhibit a single distribution of zeta potential that is highly positive (≈ 40 mV), the presence of nanomolar albumin concentrations induces the formation of liposome sub-populations with reduced zeta potentials, although still positive, as well as a sub-population with a similar charge as the liposomes in the absence of albumin (Figure S15). Hence, we suggest that insufficient protein coating enables liposome binding as there are presumably uncoated regions in the liposome surface and/or completely uncoated liposomes. Thereby, liposomes can dock and fuse with the GUVs at low albumin concentration. At intermediate concentrations, they are still able to bind to the GUVs but fusion becomes less favourable, while at higher concentrations, both docking and fusion are completely blocked. This may be due to increased surface coverage on the liposomes, shielding liposome charges and blocking docking (Figure 4 F). However, given that a sub-population of liposomes still appear to have little protein coverage at the higher albumin concentrations (Figure S15), the presence of free proteins in the environment may also contribute to the reduced liposome docking efficiency. Fusion efficiency reaches its half already at ∼ 75 nM albumin and is completely blocked with ∼ 250 nM of albumin. In summary, the results show that the presence of proteins in the dispersion blocks fusion by preventing liposome binding to the GUVs and that liposome docking and fusion seem to be separate processes controlled by the degree of corona coating that ultimately shields nanoparticle charge.

### 6. Fusion-mediated intracellular delivery of cargos in living cells and PM permeabilization

Given the rich effects of liposome fusion on model membranes, we wondered whether these effects are also mirrored in living cells, and by extension, how biophysical and biological processes are intertwined. We incubated living human embryonic kidney (HEK) cells with cationic liposomes and tested their ability to fuse, deliver encapsulated water-soluble cargos, and potential PM disruption. The experiments were performed in the absence of a protein corona, while we discuss the results for liposomes coated with a corona in the next section. Cyan labelled Atto647-PE was used as lipid marker with liposomes encapsulating Dextran 3 kDa (green) as a water-soluble cargo model, and cell incubation was carried out in the presence of SRB (red) in the outside medium to test for PM permeabilization (Figure 5 A). MVL5 liposomes exhibit weak PM fusion signal. Instead, they are predominantly internalized by cells, as observed by co-internalization with SRB (Figure S16 and Movie S2). For DOTAP liposomes, in contrast, efficient lipid and content mixing is observed (Figure 5 B) when cells are incubated with 300 μM liposomes (lipid concentration) for 30 minutes, consistent with fusion. In the early stages of incubation with the liposomes, the lipid marker is preferentially located in the plasma membrane, whereas it is internalized at later stages, although the intracellular localization of the membrane markers depends on the probe used (Figure S17). Importantly, cells do not become permeable with this vesicle concentration, despite the more than one order of magnitude higher concentrations used compared to those that were found to disrupt GUVs.

**Figure 5.**
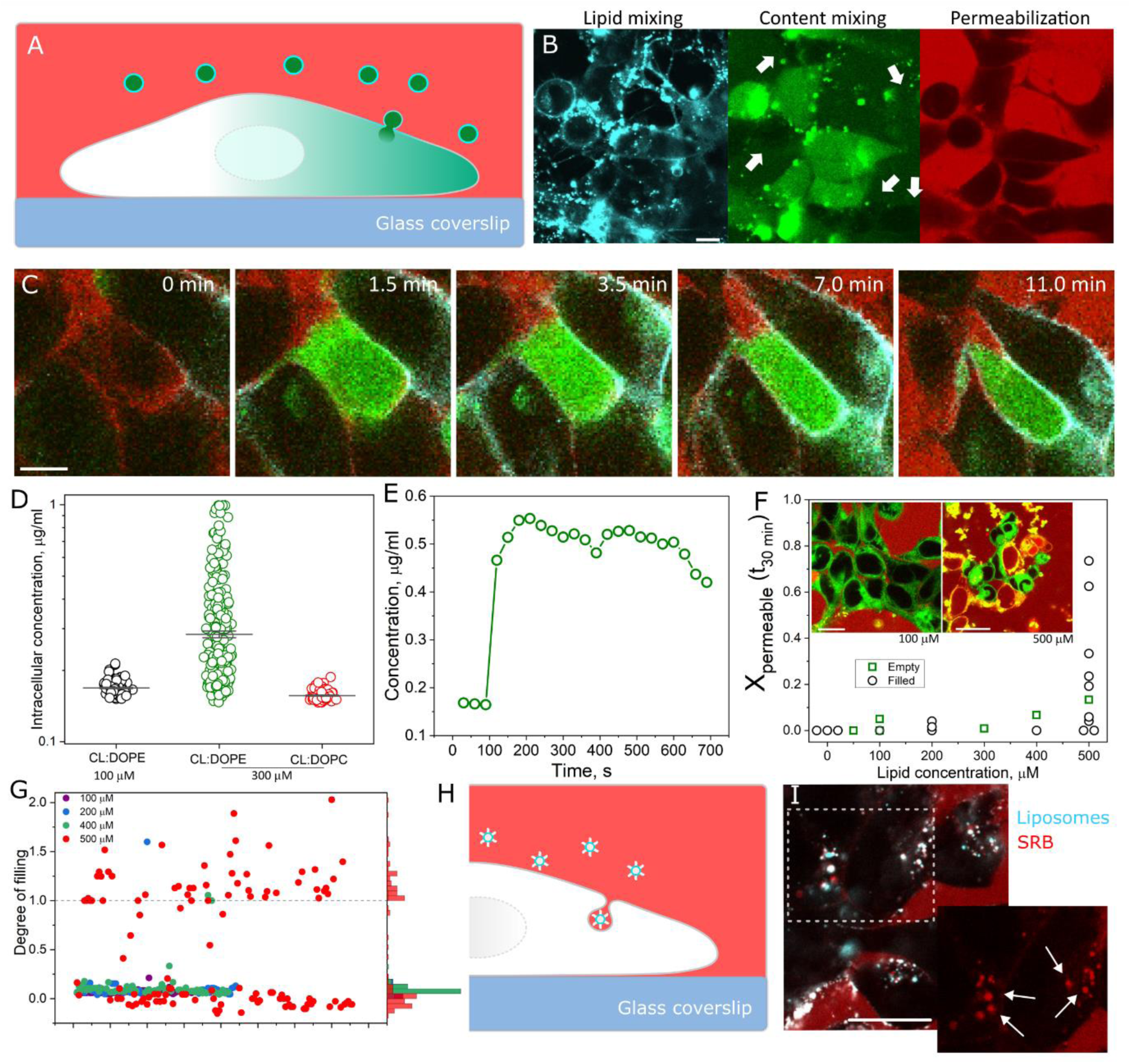
Intracellular delivery of cargos is associated with plasma membrane permeabilization only at high liposome concentration. A, sketch of the putative delivery process. Liposomes (cyan membrane) deliver the encapsulated water-soluble cargo (green) by direct fusion with the PM of living cells. B, representative confocal images of DOTAP:DOPE liposomes (1:1 mole fraction) incubated with living HEK cells at 300 μM lipid concentration. Exposure results in the transfer of Atto647-DOPE from the liposome membrane to the PM as well as the delivery of Dextran 3 kDa, green (encapsulated at a nominal concentration of 40 μg/ml). Incubation was done in the presence of 10 μM SRB as a cell leakage marker. Arrows in the Dextran channel point at cells whose Dextran 3 kDa signal is below detection sensitivity (and hence not visible in this channel). C, real-time visualization of intracellular delivery of water-soluble cargo. Note that due to its small size (∼ 1 nm) Dextran 3 kDa can locate in the nucleus. D, intracellular Dextran 3 kDa concentration for the cell shown in panel C. The decrease in intracellular Dextran signal at later times is due to the cell moving slightly out of focus. E, single cell intracellular Dextran 3 kDa concentration for different liposome compositions and concentrations after 30 minutes of incubation. Mean and standard error of the mean are shown. Note the log scale. F, fraction of cells that become permeable to SRB as a function of liposome (DOTAP:DOPE, 1:1 mole fraction) concentration (total lipid) during 30 minute incubation. Each data point represents the measured fraction of tens of individual cells. Black and green data represent empty and Dextran 3 kDa-filled liposomes. Inset, image of cells incubated with 100 μM or 500 μM DOTAP:DOPE (1:1 mole fraction) liposomes labelled with Bodipy C_16_. G, measured degree of filling of individual cells for several liposomal (total lipid) concentrations. The dashed line indicates the expected values for complete permeabilization. Values above 1 are a result of non-specific SRB affinity for intracellular structures. Histograms are also shown. H, sketch of protein-coated liposomes co-internalized with SRB by cells. I, overlay of DOTAP:DOPE:DOPC (20:50:30 mole fraction) liposomes (500 μM lipid concentration, labelled with 0.5 mol% Atto647-DOPE, cyan) and SRB after pre-incubation with 10x-diluted serum. Scale in C: 10 μm. All other scale bars: 20 μm. The inset shows the magnified image from the dashed square in panel I.

As shown in Figure 5 B (arrowheads), there is a fraction of cells whose amount of delivered cargo is below our detection sensitivity, showing that the delivery process is heterogeneous. In order to quantify the delivery, we made an, admittedly rough, estimate using the calibrated signal of free Dextran 3 kDa (Figure S18). The mean intracellular concentration of Dextran 3 kDa (when encapsulated at 40 μg/ml in DOTAP:DOPE liposomes and when the liposomes are administered at 300 μM lipid concentration) is 0.28±0.17 μg/ml (mean±s.d.), with as much as 1 μg/ml in some cells (Figure 5 D). Note that due to its small size (∼ 1 nm diameter^[53]^), the probe can also enter the cell nucleus. The maximum observed intracellular dextran concentration (1 μg/ml) corresponds to approximately 10^5^ individual liposomes fusing to a single cell. This estimate is based on an average HEK cell volume of 5000 μm^3^, approximating the cell as a 10-15 μm diameter hemisphere, and an average liposome diameter of 100 nm. Decreasing liposomal concentration to 100 μM or using DOPC as the helper lipid results in a much lower level of intracellular delivery and an even larger heterogeneity across the cells, although the amount of delivery material can still be high in some cells (Figure S19). Of note, the intracellular delivery is a very fast process. Figure 5 C and Movie S3 show the real-time delivery of Dextran 3 kDa to an individual cell, reaching ∼ 0.5 μg/ml concentration for the observed cell (Figure 5 E). Intracellular delivery of this significant amount of cargo happens within a few seconds, showing the high delivery speed. While it has been shown in GUV models that fusion of cationic liposomes takes place in a few seconds or faster^[38]^, these results show that intracellular delivery of cargos are comparatively fast in living cells, corresponding to orders of magnitude faster delivery compared to internalization-based pathways^[54–57]^.

We next tested the susceptibility of the plasma membrane to damage by incubating HEK cells with increasing liposome concentrations and measured the fraction of cells that become permeable to SRB within 30 minutes. As with GUVs, disruption is assessed by PM membrane permeabilization and SRB entry. Real-time observation is necessary since a small but detectable fraction of cells (∼ 5%) are intrinsically permeable to SRB, most likely corresponding to dead cells, and hence only those that became permeable during liposome treatment were considered for analysis. For intact cells in the absence of liposomes, SRB has no affinity for the PM nor does it diffuse in or is internalized by the cells during the observation period (Movie S4), whereas SRB binds intracellular structures in chemically-permeabilized cells (Figure S20). Upon exposure to liposomes at a 100 μM lipid concentration, extensive mixing is detectable and the cells do not change their morphology nor do their PM become permeable to SRB (Movie S5). At 400 μM, more intense mixing is detected and some cells exhibit morphological changes although with no sign of PM permeabilization (Movie S6). Only at a 500 μM lipid concentration do some HEK cells become permeable to SRB (Movie S7). As shown in the inset in Figure 5 F, permeable or intact cells are clearly identifiable. Figure 5 F shows the fraction of cells that become permeable to SRB as a function of liposome concentration. Note that although there is some variation between different experiments, significant PM permeabilization is only observed at the higher liposome concentrations. These effects are independent of whether the nanoparticles are empty or whether they are filled with cargo. Of note, in the absence of corona, PM damage only occurs upon a combination of high liposome concentration and high σ_M_, whereas 500 μM lipids but lower σ_M_ liposomes do not permeabilize cells (Figure S21).

More detailed permeability analysis at the single cell level shows that PM permeabilization is an all-or-none mechanism (Figure 5 G). The values above 1 are a result of the intrinsic SRB affinity for intracellular structures. All-or-none permeabilization indicates that the pore lifetime or pore size is longer/larger than that in GUVs, and the pores remained open until at least intracellular equilibration of SRB concentration. Notably, for permeable cells whose morphology is not or only mildly altered, the delivered Dextran 3 kDa cargo remains entrapped despite PM permeability to SRB (Figure S22). This shows that the pores formed have a size threshold, and even the formation of small pores does not necessarily forbid the use of cationic liposomes as efficient delivery vehicles, especially if the cells are able to recover the injury once the liposomes are removed from the medium (see below). While the cell density and available membrane area per cell are very different to those in the GUV experiments, the results suggest that cells are more resilient against disruption than GUVs, although this should be interpreted carefully. In summary, liposomes enables fast delivery of large amounts of cargos, although heterogeneously, and the liposomal concentration required for intracellular cargo delivery is below that which induces PM disruption.

### 7. Liposome fusion fluidizes the PM of HEK cells

From GUV model membranes, changes in fluidity are a result of the final membrane composition after fusion. Given the very ordered nature of the PM of cells^[58]^, we thus anticipate that liposome fusion, and thus transfer of DOTAP and DOPE lipids, is expected to lead to an increase in membrane fluidity, which using FliptR as a fluidity marker would result in a decrease in lifetime. FliptR strongly accumulates at the PM for extended periods of time in HEK cells (Figure 6 A and 6 B), as for other cell lines^[49]^. For control HEK cells in the absence of liposomes, the lifetime measured inside the cells (i.e. intracellular membranes) is lower compared to that of the PM (Figure 6 D and Figure S23 and Figure S28), demonstrating the PM’s high packing as previously reported^[48,49,58]^. The measured lifetime at the PM is 5.55±0.12 ns for at least 30 minutes observation (Movie S8), in accordance with other studies^[48,49]^. Interestingly, incubation with non-labelled cationic liposomes containing 20% or 50% DOTAP results in liposome labelling (due to free FliptR in the medium) and internalization, possibly along with internalization of some PM material (Figure 6 C and 6 D and Movie S9). As a result of stronger intracellular signal compared to the control, the signal from whole cell measurements, otherwise dominated by the longer lifetime of the PM, is now skewed to shorter lifetimes due to the contribution from the bright intracellular structures in a liposome concentration-dependent manner (Figure 6 G and 6 H). The decrease in whole cell lifetime is fast and occurs in less than 5 minutes though there are cell-to-cell variations.

**Figure 6.**
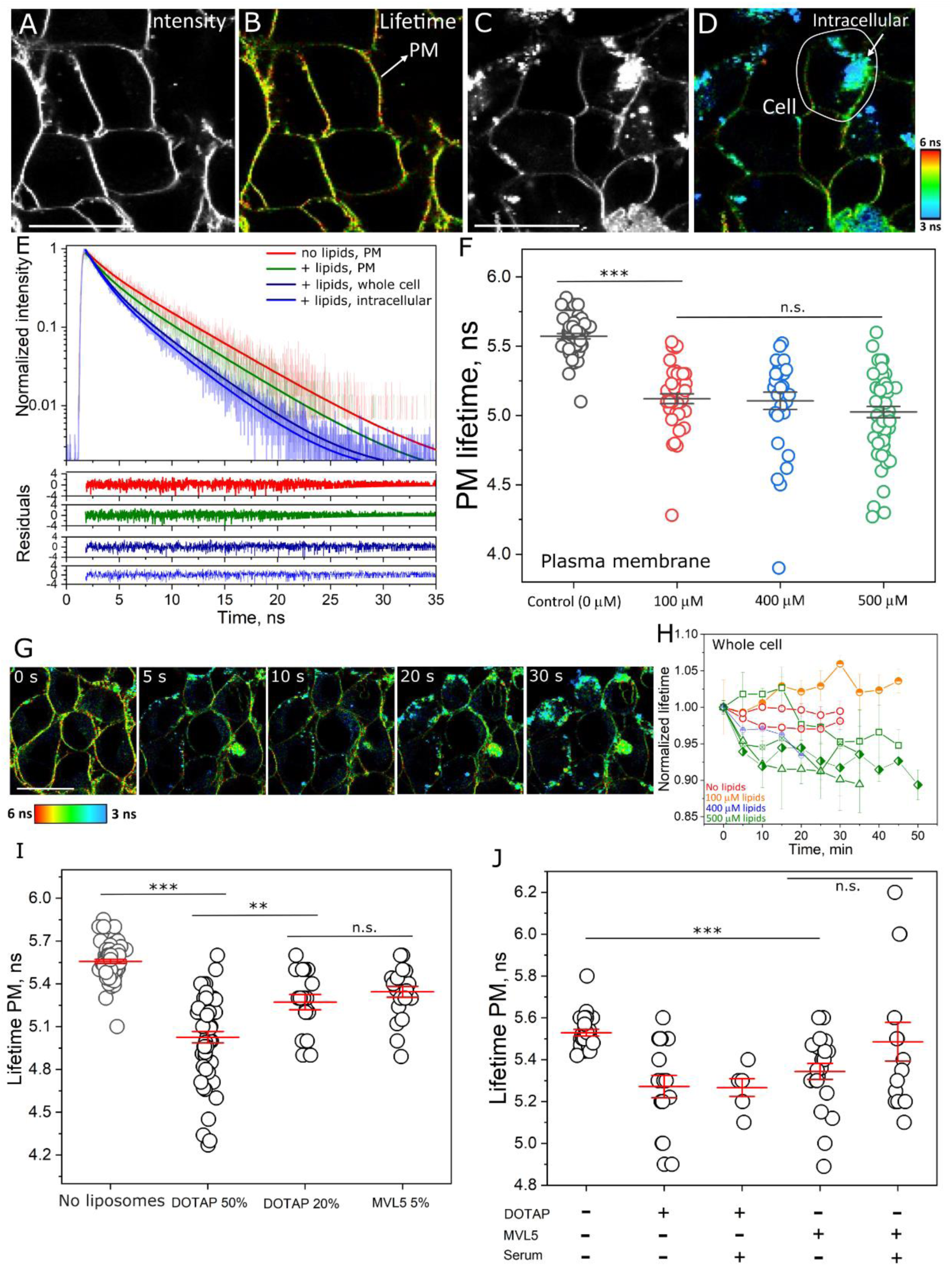
Liposome fusion fluidizes the plasma membrane. A, cells labelled with FliptR (1 μM in the cell medium) retain the probe at the PM for extended periods (at least 30 minutes) as observed from fluorescence intensity. B, FLIM image of the same cells, highlighting the PM. C, extensive bright puncta upon incubation with non-labelled 500 μM (total lipids) DOTAP:DOPE (1:1 mole fraction) liposomes. The reduced PM signal compared to control without liposomes indicates that some of the PM has been internalized. D, FLIM image of the same group of cells shows that the FliptR’s lifetime in the PM is reduced. The intracellular puncta/bound liposomes (arrow) exhibit an even shorter lifetime. E, measured fluorescence decay along with the fits and residuals from the fit for the structures shown in B and D. For all regions and systems, FliptR’s lifetime is best fitted with a double exponential decay model. F, long lifetime measured on the PM for control cells (no liposomes) and cells incubated with increasing concentrations of DOTAP:DOPE (1:1 mole fraction) liposomes. Each circle represents the measurement on a single cell. Statistical significance assessed by an independent two-sample *t*-test at a level of 0.05. G, time-lapse FLIM images of a group of cells incubated with 500 μM (total lipids) DOTAP:DOPE (1:1 mole fraction) liposomes. H, representative whole cell dynamics of lifetime changes. Errors (standard errors of the mean) indicate variations from different cells. The lines are a guide to the eye. I, measured FliptR long lifetime at the PM for liposomes containing DOTAP or MVL5 as the CL and with different σ_M_. J, effects of serum on the changes in PM lifetime. All scale bars: 20 μm.

When we measure FliptR lifetime specifically at the PM, we also observed clear PM fluidification (Figure 6 F). Again, this depends on liposome concentration (Figure 6 F and Figure S23). The extensive incorporation of external lipid material via fusion not only decreases PM packing, but it also decreases membrane tension in GUVs^[38,59]^ as well as in cells^[60]^, and thus it is tempting to speculate that the appearance of intracellular membrane material of lower fluidity may be comprised of internalized vesicles. Interestingly, PM fluidification occurs at liposomes concentrations lower than those necessary to induce pore formation and has, in fact, already levelled off at 100 μM lipid concentration. An average similar PM fluidification is observed at low σ_M_ using DOTAP or MVL5 liposomes, although with a reduced fraction of highly fluid membranes (Figure 6 I). This not only shows that the changes in fluidity also depends on σ_M_, but also indicates that these changes occur earlier than induced PM permeabilization. We conclude that the extensive incorporation of lipids in the PM due to direct liposome fusion fluidizes the PM and likely reduces surface tension, leading to internalization of PM material, in a way that depends on liposome concentration and charge density, and that changes in fluidity are milder effects compared to PM pore formation.

### 8. Effects of protein coating on liposome-cell interactions, fluidification and PM permeabilization

Pre-incubation with serum screens the charge and, in general, interactions of cationic liposomes, most likely by completely coating their surface, as also observed for other particle models^[43,61]^, preventing fusion with GUVs. We next tested whether corona formation also prevents particle binding and fusion with living cells. Here, cells were exposed to the liposomes in the presence of both a hard and soft corona, as well as in the presence of serum proteins in the medium, comparable to the experiments presented in Figure 4. Liposome internalization and content mixing can be easily distinguished by their staining pattern; whereas content mixing leads to homogeneous labelling of the PM with the lipid marker as sketched in Figure 5 A and shown in Figure 5 B, internalization leads to a spotted pattern. Furthermore, as vesicles formed during the internalization process inevitably engulf some extracellular liquid content, SRB becomes co-internalized with liposomes and thus internalization is detected as cyan-red spotted pattern as sketched in Figure 5 H and shown in Figure 5 I. Figure S24, Movie S10 and Movie S11 show that DOTAP and MVL5 liposomes upon pre-incubation with serum are internalized by cells rather than directly fusing to the PM. Of note, these liposomes do not induce PM permeabilization, even at the highest concentration tested (Figure S25) although they also fluidize the PM, regardless of the CL used (Figure 6 J and Figure S26). Nevertheless, the fraction of cells with highly fluid PM is reduced, showing that the effects of protein-coated liposomes on membrane properties are milder. In summary, protein corona formation does not prevent liposome binding to the PM of living cells but it does change the mode of interaction, switching from content mixing to internalization. This is reflected in their much milder mechanical effects on cells.

### 9. Changes in material properties are transient

All experiments above involve the assessment of GUVs and cells upon acute treatment with liposomes. For drug and gene delivery applications, many protocols involve a short pulse with the delivery vehicle, followed by vehicle removal to reduce potential side effects. If the vehicle induces toxic effects, as is typical for cationic particles^[18,39,40,43]^, particle removal may allow surviving cells to recover whilst still achieving cargo delivery. To test whether the effects observed in our system, namely pore formation and changes in membrane fluidity, are permanent or temporary, we incubated the cells with increasing concentrations of liposomes as above, removed the liposomes after 30 minute incubation, and placed the cells back under cell culture conditions for 24 h to allow cell recovery. When compared to the control cells (treated with sucrose solution in the absence of liposomes), the liposomes induce a similar fraction of dead cells (∼ 20-40 %) for all concentrations tested (Figure S27). Note that the fraction of dead cells is already higher in sucrose solution itself compared to cells with cell medium alone, but sucrose (or other low ionic strength solution) was required due to liposome preparation and for better comparison with the GUV experiments. After the recovery period, the remaining cells (∼60-80 %) were incubated with SRB to assess whether PM pores were still present. No intracellular signal of SRB was observed, indicating that the PM was intact, even at the liposomal concentrations that induced significant PM damage (Figure S28 A and Figure S28 B). As can be seen in Figure S28 C, FliptR, which was added to the cells before imaging (after the recovery period), largely stains the plasma membrane for all concentrations tested, with intracellular labelling indistinguishable from that of control conditions without liposomes. This indicates that incorporated lipids, internalized PM materials and liposomes have been fully processed. Importantly, FliptR’s lifetime measured at the PM is identical in all conditions, even at concentrations well above those that induced changes in acute treatment (Figure S28 D). In summary, while administered liposomes are able to deliver cargos into living cells, any measured adverse effects on PM membrane integrity, changes in membrane fluidity or presence of large amounts of internalized lipid materials are recovered to steady conditions upon liposomal removal.

## Discussion

In this work, we show that cationic fusogenic liposome nanoparticles efficiently deliver encapsulated cargos to GUV model membranes and living cells, though they modify a number of fundamental material properties of the target membranes in the process. Figure 7 summarizes these observations. Fusion increases membrane area upon incorporation of new lipids, initially decreasing membrane tension, as observed in GUVs by the appearance of membrane fluctuations and increase in volume, and as indicated in cells by the internalization of PM material. Depending on the initial membrane composition, the newly added lipids may increase or decrease membrane packing and fluidity, which tend towards values intermediate between those of the acceptor membrane and liposomes. In cells, fusion fluidizes the PM. Due to their small sizes, liposomes carry an intrinsic leaflet asymmetry, with more lipids in their outer than inner leaflet^[62]^. Fusion of a large enough number of liposomes brings this asymmetry to the acceptor membranes, leading to budding as well as an increase in curvature tension that overcomes the initial decrease in tension from the gained area. In extreme conditions, curvature tension build-up results in the formation of membrane pores of specific sizes, and they may retain the delivered cargo depending on cargo size. Interestingly, the formed pores are transient and they close in a few minutes in GUVs. For corona-coated liposomes, charge shielding prevents strong electrostatic association with the acceptor membranes, hindering binding and fusion with GUVs, and shifting the mechanisms of interactions with cells from direct fusion to internalization. While these modifications do not necessarily mean serious adverse effects, because under certain conditions the cells are able to fully recover to pre-liposome treatment state, they are worth considering when designing a drug delivery vehicle. To the best of our knowledge, drug delivery-mediated changes in the material properties of synthetic and biological membranes were until now unknown.

**Figure 7.**
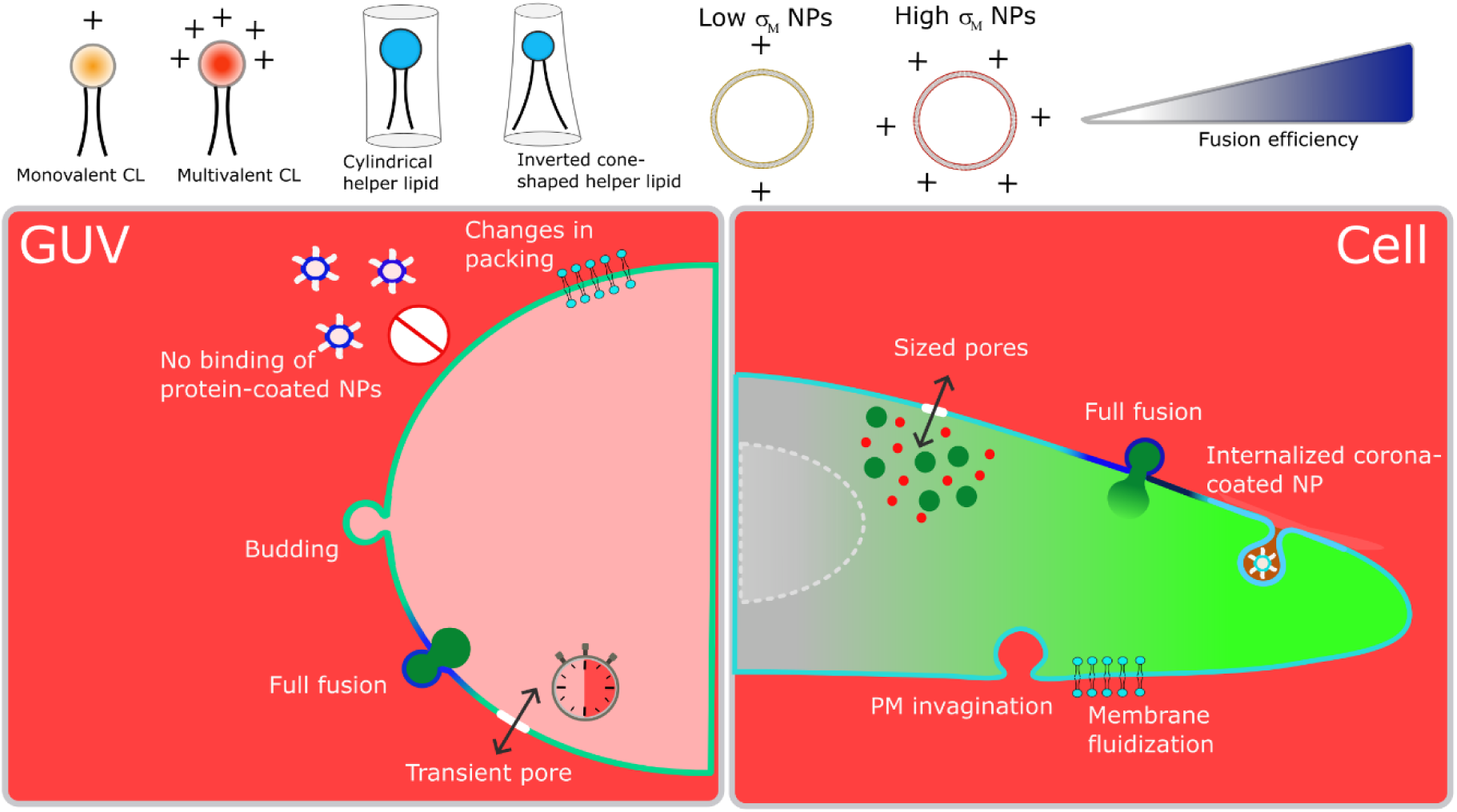
A summary of fusogenic liposome nanoparticle interaction with model membranes and cells. Top left: monovalent (orange headgroup) or multivalent (red headgroup) cationic lipids (CL) and cylindrical or inverted cone-shape helper lipids. Top right: liposome membrane charge density can be tuned by increasing the mole fraction of CL. Bottom panels: main observations of changes in the material properties of GUV membrane models and living cells. Bottom left: in GUVs, fusion occurs via (i) content mixing, leads to (ii) changes in membrane fluidity/packing, which can (iii) increase/decrease depending on the initial and final membrane compositions, (iv) leads to bud formation due to changes in spontaneous curvature and (v) the opening of transient hydrophilic pores. Heavily corona-coated liposomes are unable to bind. The results are independent of the chemical composition of the CL or helper lipid. Bottom right: in cells, cationic liposomes directly fuse with the PM, leading to a fast and efficient delivery of cargos in the cytosol. Fusion fluidizes the membrane as external low T_m_ lipids are incorporated in the highly packed PM. Extensive incorporation of lipids likely leads to active internalization of PM content to counterbalance a decreased tension. Fusion also leads to the formation of pores of defined sizes that allow the entry of small probes from the medium but the delivered large cargos are retained. Cationic liposomes heavily coated with a protein corona are not able to directly fuse with the PM but efficiently bind to the membrane and are internalized.

### Effects of charge density and helper lipid

In many ways, intracellular delivery of encapsulated water-soluble cargos via direct PM fusion mirrors the activity of cationic liposomes used in gene delivery. For lipoplex nanovehicles, high σ_M_ leads to efficient gene delivery independent of the helper lipid, although the helper lipid determines the interaction mechanism; from complex internalization with cylindrical lipids to direct fusion with the PM with inverted cone-shape helper lipids^[32,63]^. Similarly to transfection of foreign genes in cells^[63–65]^, or the interaction of other types of charged nanoparticles^[66]^, binding and fusion of cationic liposome nanovehicles with acceptor synthetic and biological membranes depends on σ_M_. At high σ_M_, liposome binding is almost instantly followed by fusion and the helper lipids are dispensable, whereas at intermediate or low σ_M_, membrane-membrane interaction is weakened and fusion is delayed and inverted cone-shape lipids enhances fusion. Similarly, bound weakly charged nanoparticles may eventually diffuse away upon binding^[66]^. Delayed fusion gives the cells the ability to internalize the bound particles as we have recently shown^[57]^. In contrast, non-charged liposomes or liposomes whose charges have been completely screened by a protein corona coat are unable to bind to model membranes although they do bind to cells. In this case, binding results in internalization rather than direct fusion. For small solid cationic nanoparticles, both Chol in homogeneous membranes or defect regions induced by membrane phase separation increase binding and lipid exchange (the equivalent of fusion in our system) to acceptor anionic membranes^[66]^. For cationic fusogenic liposomes, Chol does not seem to increase binding, although it does decrease fusion due to an increase in membrane stiffness^[36]^, and does modify the interaction energetics^[67]^. Furthermore, the presence of membrane defects also results in higher liposome fusion efficiency^[68]^. High enough σ_M_ bypasses the need of these factors. Hence, despite the extremely diverse nanoparticle type and delivery mechanisms, the effects of charge density in drug delivery vehicles seems to be more general than might have been anticipated.

### Nanoparticle toxicity

Cationic lipids do not exist in nature and administration of cationic particles, including liposomes, has shown to induce toxicity via various mechanisms^[69–71]^. One of such proposed mechanisms includes the need for the cell to metabolize these foreign species^[63,72]^. The use of multivalent lipids could potentially circumvent nanoparticle toxicity by achieving high σ_M_ at low CL fractions, reducing metabolic demands on the cell^[72,73]^. However, as we showed for both synthetic and biological membranes, for comparable amounts of administered nanoparticles, σ_M_ also determines toxicity (as assessed from PM permeabilization) irrespective of the chemical composition of the CL. To reduce toxicity, one strategy could include nanoparticle coating with inert proteins or polymers. However, coating not only reduces fusion efficiency at moderate surface coverages, creating a charge/coating dilemma similar to the PEG dilemma observed with stealth liposomes^[8,74]^, but also shifts the interaction to particle internalization. A more reasonable alternative may constitute the use of cleavable components as shown for PEG-coated liposomes^[65]^, in which on-demand removal of the coating would expose nanoparticle charge, and hence induce fusion, although this remains to be demonstrated. Importantly, membrane leakage occurs at liposomal concentrations that are higher than that required to achieve fusion and cargo delivery and thus, for a given σ_M_, there is a relatively large range of liposome concentrations in which intracellular delivery is efficient and leakage is minimum, although the PM does fluidize. These disruption effects mirror the ability of RNA delivery to cells mediated by lipoplex nanoparticles, with gene silencing below the onset of cytotoxicity ^[64]^. Furthermore, the formed pores have a size threshold, and larger delivered cargos (here above ∼ 1 nm, the expected hydrodynamic radius of Dextran 3 kDa^[53]^) are retained in the cell even if the PM is permeabilized. Lastly and most importantly, upon liposomal removal, the cells are able to fully recover to their state before contact with liposomes.

Taking a very simplified interpretation of the observed results by treating them as purely physical in nature, they can be understood by the mechanical effects of fusion of a large number of liposomes to acceptor membranes. Membrane pore dynamics are controlled by a balance between surface tension and edge tension, favouring pore opening and closure, respectively^[75]^. In typical phospholipid bilayer membrane models, edge tension is high and mechanically opened pores spontaneously close^[76,77]^, but a decrease in edge tension will favour the open pore state^[45,78]^. In stably open pores, SRB leaks in to full equilibration in ∼ 5 minutes^[45]^. As in the experiments here leakage was incomplete, we assume that the pores close in the first minutes after pore opening, and thus fusion-dependent decrease in edge tension is significant, but not strong enough to lead to GUV collapse. The discussion above is nonetheless valid for symmetrical artificial membranes. Due to the intrinsic leaflet number asymmetry in small liposomes, their membranes have more lipids in their outer than their inner leaflets. Fusion of large enough numbers of liposomes is thus expected to also result in asymmetry in the GUVs, consequently inducing GUV bud formation outwards as observed. Asymmetrical distribution of lipids across the leaflets reduces the edge tension^[79]^, favouring the state of longer-lived pores. A recent study showed that the number of lipids across the PM leaflets in living cells is also asymmetrically distributed, which poses consequences for PM mechanical stability^[80]^. We are not aware of edge tension values in cells, but extensive liposome fusion may also introduce or even increase such asymmetry. It is thus tempting to speculate that, similarly to GUVs, membrane asymmetry and a decrease in edge tension may also be the reason why pores are formed in cells upon liposome fusion. If such effects take place more quickly than the ability of cells to cope with the introduced stress, pores will likely open in the PM, as indeed observed. On the other hand, if the cells survive treatment, removal of liposomes ceases stress build-up and cellular activity may ensue to counterbalance the effects of newly introduced foreign lipids, recovering membrane fluidity, and activate membrane flippases and scramblases, relaxing leaflet imbalances.

### GUVs as cellular membrane models

To what extent can GUVs or other model membranes be a predictor of the behaviour of biological membranes and cells? GUV membrane models, while not bearing all cellular components, can reconstitute the fundamental processes at the molecular level with incredibly high robustness. A great advantage includes the easy tuning of their material properties by the appropriate choice of their components. In the specific case of cationic liposomes, particle binding, fusion and cargo delivery, pore formation and changes in overall material properties, such as fluidity and curvature, that were observed in GUVs were mirrored in live cell experiments. However, despite being an excellent tool, GUVs do not always predict all cellular outcomes, and even the (nearly) identical outcomes occur at vastly different liposome concentrations. On the one hand, membrane fluidity changes can be directly inferred from the changes in lipid composition, qualitatively observed in both GUVs and cells. On the other hand, major differences observed between GUVs and cells included the interaction of protein-coated liposomes with cells, leading to nanoparticle internalization by cells, whereas they were completely unable to bind GUVs. The switch to cellular internalization mechanisms for protein-coated liposomes likely stems from the specific binding of biomolecular corona-proteins to cell membrane receptors, which consequently triggers their endocytosis. Indeed, the internalization pathway of several liposomal formulations, such as Doxil, has been shown to be mediated by the presence of particular proteins in the corona^[81–83]^. Furthermore, whereas liposome fusion induces spontaneous curvature on GUVs, resulting in outward budding formation, in cells, intracellular vesicles and PM internalization were observed. While in GUVs, budding is a purely physical process that is associated with the asymmetric incorporation of lipids from the intrinsically asymmetric liposomes, resulting in more lipids in the outer than the inner leaflets^[38]^, in cells, PM internalization is most likely actively driven. The sudden incorporation of excess of lipids via fusion acutely decreases PM tension activating the endocytosis machinery for lipid recycling^[84–86]^. Furthermore, the high compositional complexity and interaction network of the PM with other cellular factors (i.e. cytoskeleton^[87–89]^, media with varying ionic strengths^[45,90]^, PM asymmetry^[34,80]^, and the presence of protein machinery responsible for resealing pore damage ^[91]^) likely reflects the higher stability of cells compared to free-standing GUV membranes.

### Direct membrane fusion versus internalization-based pathways

Most nanoparticle-based drug delivery strategies are, one way or another, based on particle internalization by cells, with a few exceptions^[92,93]^. This means that, at least transiently, the particles are entrapped in intravesicular compartments, such as endosomes and lysosomes. However, internalization alone is not sufficient for delivery since the vehicle-cargo complexes are still unavailable. In fact, corona-coated liposomes that were internalized by cells were unable to release the encapsulated cargo during the observation time of our experiments, for any CL or helper lipid used and at any concentration. Hence, the cargo, either when complexed with its vehicle or (preferably) alone needs to be released from these compartments. In general, vesicular processing is slow and can take several hours^[94]^. Long processing times in low pH and in the presence of proteases and nucleases increase the chance of cargo degradation. Thus, even in the case of successful but delayed release, the cargo might no longer be functional. Strategies to improve endosomal escape efficiency and speed includes osmotic endosomal inflation using internalized charged polymers that induce ion and water pumping into the vesicles via the proton-sponge effect^[95,96]^. This nevertheless not only results in cytosolic release of proteases, but the number of ions needed to burst endosomes has been suggested to increase quadratically with endosomal radius, limiting escape to small endosomes^[97]^. As shown in cells, although siRNA in complex with cationic liposomes can be efficiently internalized, release from endosomes through endosomal pores is transient and incomplete^[98]^. In fact, endosomal escape is a more limiting factor in drug/gene availability than the degree of internalization^[97]^, with as little as 1-2% of internalized cargo being effectively released from the endosomes^[99]^. Due to their presumably high (but unknown) edge tension, formed pores in the endosomal membranes are likely short lived, limiting escape to small cargos and only while the pore is open, probably explaining the extremely low escape and incompleteness of cargo release from internalized vesicles.

The highly fusogenic liposomes undergo fusion with the first contacting membrane (i.e. the PM and the contacting membrane of the GUVs) within a few seconds, faster than most internalization-based pathways^[100,101]^, with direct and immediate delivery of large amounts of cargos in the cell cytosol. This bypasses slow and inefficient internalization pathways, with the additional advantage that neither the vehicle nor the cargo are exposed to the low pH and enzymatic milieu of intracellular vesicle compartments. Direct membrane fusion also circumvents the need for complex surface functionalization and intracellular processing, such as the use of pH-sensitive moieties^[102,103]^ or polymers that aid endosomal escape^[104,105]^, significantly simplifying formulation design. Direct fusion-based intracellular delivery is very likely extendable to any water-soluble cargo and amenable to cargo co-delivery^[106]^, and it is not limited by cargo size or specific chemical identity, as successfully demonstrated for proteins^[107,108]^, oligonucleotides^[109]^ and nanoparticles^[108,110]^. The many advantages of direct fusion-based cargo delivery using fusogenic liposomes make them extremely attractive nanovehicles in drug delivery. The discovery that the extensive fusion of a large number of liposomes transiently modifies the material properties of acceptor membranes and cells in their course of cargo delivery will help the development of more efficient delivery systems with limited and controlled side effects.

## Conclusions

Drug delivery nanoparticles are excellent tools to increase the availability of cargos at sufficient amounts in the target location, ideally quickly, while being inert; that is, they should not induce cellular responses or alter the cell state upon cargo delivery. Here, we show that highly charged cationic liposomes fuse with synthetic vesicles and the biological PM of living cells to efficiently deliver large amounts of encapsulated cargos within seconds, bypassing slow and often inefficient internalization-based pathways. However, in the course of cargo delivery, they modify several material properties of the acceptor membranes, including membrane packing, spontaneous curvature and surface tension and they lead to the formation of transient pores of defined sizes. To the best of our knowledge, this is the first study to systematically assess the effects of a drug delivery vehicle on the mechanical properties of membranes. We show that for cationic liposomes, the molecular mechanisms of toxicity are associated with a combination of local changes in composition, curvature and tension. Importantly, there is a relatively wide concentration range that achieves effective intracellular delivery, and although some cell toxicity is observed, this is reversed by removal of liposomes after successful delivery. These effects depend on liposome charge density and can be further modulated by the type of helper lipid. Due to their high charge, intracellular delivery is limited to *ex vivo* applications since in the body serum proteins would quickly bind and cover their surface. However, they could be fairly adaptable to *ex vivo* applications with efficient and fast administration in serum-free media and with limited side effects once the cells return to physiological conditions. In many aspects, fusion-based drug delivery mirrors lipoplex systems despite major differences in structures and interactions with the target membrane, indicating a more universal role of charge than anticipated. We expect the results to aid the design of more efficient drug delivery systems with limited toxicity.

## Materials and methods

All chemicals and materials were used as obtained. The phospholipids 1,2-dioleoyl-sn-glycero-3-phosphocholine (DOPC), 1,2-dioleoyl-sn-glycero-3-phospho-(1’-rac-glycerol) (sodium salt) (DOPG), 1,2-dioleoyl-sn-glycero-3-phosphoethanolamine (DOPE), L-α-phosphatidylserine (Brain, Porcine) (sodium salt) (PS), 1,2-dioleoyl-3-trimethylammonium-propane (DOTAP), N1-[2-((1S)-1-[(3-aminopropyl)amino]-4-[di(3-amino-propyl)amino]butylcarboxamido)ethyl]-3,4-di[oleyloxy]-benzamide (MVL5) and cholesterol (plant derived), as well as the fluorescent dye 1,2-dipalmitoyl-sn-glycero-3-phosphoethanolamine-N-(lissamine rhodamine B sulfonyl) (ammonium salt) (DPPE-Rh) were purchased from Avanti Polar Lipids (Alabaster, AL). DOPE-Atto647N was purchased from AttoTech (Siegen, Germany). Lipid solutions were prepared in chloroform and stored at – 20 °C until use. Glucose, sucrose, Dimethyl sulfoxide (DMSO) and the fluorescent probes Bodipy C_16_ (BODIPY™ FL C16; 4,4-Difluoro-5,7-Dimethyl-4-Bora-3a,4a-Diaza-s-Indacene-3-Hexadecanoic Acid), sulforhodamine B (SRB), Dextran-FITC 3 kDa, Albumin–fluorescein isothiocyanate conjugate (FITC-albumin) were purchased from Sigma-Aldrich (St. Louis, MO, USA). Bovine serum albumin (BSA) and Poly(vinyl alcohol), PVA 87-90% hydrolyzed were also obtained from Sigma-Aldrich. The molecular sensor FliptR plasma membrane (commercially known as Flipper-TR®) was purchased from Cytoskeleton inc. (Denver, CO, USA).

### Vesicle preparation

GUVs were prepared using the PVA method^[111]^ with minor modifications^[112]^. Approximately 100 μL of a 2% (weight/volume) PVA solution (in water) was spread on two glass coverslips and the water was left to evaporate by placing the coverslip on a hot plate at ∼ 60°C (typically 10 minutes) to form a PVA film. Next, a ∼ 10 μL lipid solution (2 mM in chloroform) containing the desired lipid mixture was spread on the PVA film. Chloroform was evaporated under a stream of Argon. The coverslips were sandwiched using a Teflon spacer forming a ∼ 2 mL chamber sealed with the help of clips. The chamber was filled with a 200 mM sucrose solution (unless stated otherwise) for ∼ 30 minutes to allow GUV growth. Unless stated otherwise, growth was performed at room temperature (R.T = 18±1 °C). Hydration was carried out in the dark for lipid mixtures containing fluorescent lipids. After hydration, the GUV solution was harvested by gentle pipetting and the GUVs were used within 1 day. For imaging, the GUVs were diluted in an isotonic glucose solution to help sediment the vesicles to the bottom of the imaging chamber. If not mentioned otherwise, the GUVs were labelled with 0.5 mol% fluorescent labels, the composition of which depended on the experiment.

Liposomes made of CL:DOPE:DOPC (X:50:50-X) or CL:DOPC (X:100-X), mole fraction, in which CL is the cationic lipid DOTAP or MVL5 were prepared using the hydration-sonication method^[110]^. For the FRET experiments, the liposomes were labelled with 2 mol% DPPE-Rh. For the multicolour confocal experiments, the liposomes were labelled with 0.5 mol% DOPE-Atto647N. In short, the appropriate lipid mixture (in chloroform) was added to the bottom of a glass chamber and evaporated under a stream of Argon and further evaporated under vacuum for 1-2 hours to remove any trace of chloroform. Afterwards, the lipid films were hydrated with a 200 mM sucrose solution and vigorously vortexed until full lipid film detachment forming multilamellar vesicles (MVLs). For the encapsulation of water-soluble probes, the reporter probe was pre-diluted in the hydrating sucrose solution. SRB was added at 50 μM and Dextran-FITC was added at 0.1 mg/mL hydrating concentrations. If not stated otherwise, the final lipid concentration was 2 mM. The MVLs were sonicated using a bath sonication for ∼ 20 minutes and used within 2-3 days. GUV incubation with the liposomes was done by diluting the liposomes in sucrose to a 100 μM lipid concentration. An equivalent amount of the pre-diluted liposomes solution was mixed with 50 μL GUVs in glucose at the desired final liposomes concentration (typically 0-30 μM) for a final 100 μL solution. GUVs and liposomes were incubated for 10-15 minutes in an Eppendorf tube, after which the solution was ready for imaging. In the serum experiments for both GUVs and cells, the desired liposome concentration was pre-incubated with 10x diluted serum (or with the desired concentration of FITC-albumin in the GUV experiments) in sucrose (200 mM and 300 mM for the GUV or cell experiments, respectively) for 10 minutes, before incubation with the GUVs or cells. For imaging, the incubated samples were moved to a BSA-coated glass (1 wt% BSA) coverslip.

### Zeta potential measurements

The zeta potential measurements of the liposomal dispersions in the presence of different concentrations of FITC-albumin were characterized using a Malvern ZetaSizer Nano ZS (Malvern Instruments) and ZetaSizer Software version 7.13 (Malvern Instruments). Dispersions were prepared as above. Measurements were performed in triplicate with minimally 10 runs each at room temperature. The reported results are the mean and standard deviation across repeats.

### Cell handling, labelling and incubation

Human embryonic kidney (HEK) cells (American type culture collection, no. CRL-1573, lot no. 63966486) were used for the cell experiments. The following materials were used for cell culture and experiments: Dulbecco’s minimal essential medium (DMEM, Giboc), foetal bovine serum (Gibco), phosphate buffered saline (Gibco), trypsin (0.05 %, Gibco), trypan blue (0.4 %, Sigma-Aldrich), µ-slide 8 well chambers (Ibidi), 35 mm petri dishes (Grenier Bio-One), and a Neubauer hemacytometer (Brand). HEK cells were cultured in DMEM supplemented with 10 % serum (complete medium) at 37 °C under 5 % CO_2_ and a humidified atmosphere. All results are from cell cultures that tested negative for mycoplasma. Cells were seeded into µ-slide 8 well dishes one day prior to experiments. For multicolour confocal imaging, the cells were moved to the microscope stage at 37 °C and the cells were washed once with a sucrose 300 mM solution (isotonic to culture media). In the control permeability experiment (no liposomes), the culture medium contained 10 μM SRB. For incubation with liposomes, a liposome solution (containing 10 μM SRB) in sucrose 300 mM contained the desired final lipid concentration and pre-warmed at 37 °C, was added to the cells and imaging started immediately. As a positive control for PM permeabilization, 70 % ethanol was applied to cells for 10 min. The ethanol was then removed and replaced with SRB solution. For FLIM experiments, the cells were pre-labelled for 10-15 minutes with FliptR by adding a small volume of a concentrated FliptR solution (in DMSO) to the cell culture medium to a final 1 μM final concentration, followed by 1-2 washes in 300 mM sucrose to remove the excess of probe as well as the culture medium. Afterwards, incubation with liposomes was performed in an identical manner as above.

### Cell counting and TB staining

For the recovery experiments upon incubation with liposomes, cells were seeded into µ-slide 8 well dishes or petri dishes one day prior to the experiments. Cells were washed with warmed sucrose solution (300 mM) and then exposed to liposomes following the procedure above and then returned to cell culture conditions. After 30 min exposure, the liposomal dispersion was removed and the cells were washed with warmed sucrose solution followed by complete medium to remove any unbound particles. The cells were then returned to cell culture conditions with fresh complete medium for 24 h. The cells seeded in µ-slide 8 well dishes were then used for imaging and stained using the procedure above. To determine cytotoxicity, the cells in petri dishes were removed from the incubator after 24 h, washed with phosphate buffered saline and then trypsinized to detach them from the cell surface. The cell suspension was diluted with complete medium from which 10 µl of cell suspension was mixed with equal amounts of trypan blue and loaded into one half of a Neubauer hemacytometer. Cells were manually counted separately for four of the grid quadrants. Cells without trypan blue staining were considered to be live, whereas dead cells exhibited staining. The measurements were performed in triplicate for each liposomal concentration investigated. The reported values are the mean and standard deviation across all replicates (*i.e.* 4 quadrants per triplicate so across 12 measurements in total).

### Microscopy imaging

Multicolour confocal microscopy imaging was performed on a Zeiss LSM 710 scanning confocal microscope. A water-immersion C-Apochromat 40X/1.20 W Korr M27 objective was used for imaging. The spatial and temporal resolutions used were adjusted according to the sample conditions, but in general a consistent imaging size of 212.55µm x 212.55µm (1024 × 1024 pixels) was used in a frame scanning mode in a singular direction, with a pixel size of 0.21 µm. Line averaging = 4 and bit depth = 8. The green dyes Bodipy C_16_, FITC-Albumin and Dextran-FITC 3 kDa were excited using a 488 nm argon laser, and emission was detected between 495-555 nm. The orange dyes DPPE-Rh and SRB were excited with a 543 nm HeNe laser line and emission was detected in the range between 555-600 nm. The far-red dye Atto-647N was excited with a 633 nm HeNe laser line and its emission was detected in the range between 640-800 nm. Images were scanned in the sequential mode to minimize crosstalk between channels. Dextran 3 kDa concentration in the cells was calculated from a calibration curve. Known concentrations of Dextran 3 kDa were prepared in sucrose and their intensities were measured, exhibiting a linear response. Considering the probe intensity does not change in the cellular milieu, cells were imaged with identical imaging conditions and the intracellular concentration was directly obtained from the calibration.

The fluorescence lifetime imaging microscopy (FLIM) experiments were carried out on an inverted microscope (Olympus IX73) equipped with time correlated single photon counting (TCSPC) (PicoQuant). The samples were imaged through a 100x (1.4 NA) oil immersion objective (UPLSAPO, Olympus). Bodipy C_16_ was excited using a 481 nm laser and its emission was collected using a 525/50 nm band pass filter. The images were acquired using the SymPhoTime 64 software and all samples were excited with a pulsed 20 MHz repetition rate. Unless stated otherwise, the samples were imaged with a 128×128 pixels, 1 ms dwell-time and ∼ 300 μm/pixels with typical acquisition times of ∼ 27 s. For analysis, the GUV membrane signal at the equator or a defined region of interest (ROI) in the cell were manually selected for analysis and all pixels binned for fitting. Bodipy C_16_ fluorescence decays were fitted using a n-exponential tail fit with a single (in the absence of an acceptor) or bi-exponential decay model (in the presence of the DPPE-Rh FRET acceptor)

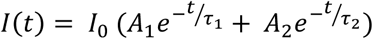

where *I*(*t*) is the intensity at time *t* and I_0_ is the intensity at *t* = 0. *A*_1_ and *A*_2_ are pre-exponentials factors associated with lifetime components *τ*_1_ and *τ*_2_, respectively^[37]^.

The amplitude-weighted mean fluorescence lifetime in the presence of the FRET acceptor (*τ*) could be calculated as

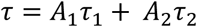

To measure the FRET efficiency (E_FRET)_, *τ* was obtained from GUVs both before (*τ*_*before*_) and after (*τ*_*after*_) incubation with the liposomes, in which FLIM-FRET efficiency is given as

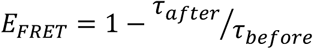

In the case of cells labelled with FliptR, a bi-exponential decay model was used from which the long lifetime τ_2_ was used for analysis. For the temperature-controlled experiments at the FLIM microscope setup, a PeCon GmbH TempController 2000-1 temperature-controller was connected to a Heating Insert P Lab-Tek™ S heating stage chamber, into which the observation chamber was placed. T-controlled experiments at the multicolour confocal microscope setup was performed in an in-built temperature-control box and controlled using the microscope software.

### Viscosity calculation from lipid diffusion

The viscosity of the GUV membranes (η_m_) was calculated from reported diffusion coefficient (D) data for the experimental systems used here^[38]^, using the Saffman-Delbrück model^[46]^

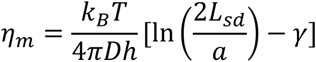

where k_B_ is Boltzmann’s constant, T is the absolute temperature, h represents membrane thickness (3.98 nm, as measured for POPC at 20°C^[113]^), *a* is the radius of the diffusing particle (assumed here as 0.5 nm) and γ ≈ 0.577, and L_sd_ is the Saffman-Delbrück diffusion length

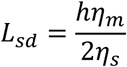

in which η_s_ is the viscosity of the surrounding fluid (1.133 cP)^[45]^. Although the model is best suited for relatively large particles embedded in or attached to the membranes (i.e. proteins) than for molecules of similar dimensions to the membrane lipid constituents (e.g. small fluorescent probes)^[114]^, it is still useful and widely used to gain information about membrane structure and properties^[115]^. Hence, regardless of the exact obtained numerical values from the estimates of membrane viscosity, the trend will be preserved.

## Author contribution

RBL designed the study, performed experiments, analysed data and wrote the first version of the manuscript. CJR performed part of the cell experiments and aided in the design of the cell experiments. JH, YBK and TJ performed part of the GUV experiments and analysed data. CÅ supervised the cell work. WHR supervised the study. RBL, CJR, CÅ, WHR contributed to the writing of the final version of the manuscript.

## Acknowledgements

We acknowledge Anna Salvati of the Groningen Research Institute of Pharmacy (Rijksuniversiteit Groningen) for providing the cells. Likewise, for access to the Malvern ZetaSizer Nano and software, we thank H. C. van der Mei and H. J. Kaper at the Department of Biomedical Engineering, University Medical Center Groningen. We thank Michiel Punter (Rijksuniversiteit Groningen) for providing the ImageJ plugin for single particle intensity analysis.

